# Clathrin-coated endocytic tubules mediate rapid and selective protein absorption by lysosome-rich enterocytes

**DOI:** 10.64898/2026.10.22.761028

**Authors:** Indrasen Magre, Tylor R. Lewis, Claudia Carugati, Carina L. Block, Siyao Wang, Kevin Li, Jieun Park, Vadim Y. Arshavsky, Daniel S. Levic, Michel Bagnat

## Abstract

In the digestive tract of neonatal mammals and in fish, lysosome-rich enterocytes (LREs) internalize dietary proteins and digest them intracellularly within a large lysosomal vacuole. Impaired LRE function leads to growth stunting and poor survival in zebrafish and mice. Previous work has shown that protein uptake relies on a scavenger receptor complex. However, how LREs can internalize proteins rapidly enough to support organismal growth is unclear. Here, using live and electron microscopy and genetic manipulations in zebrafish, we find that LREs contain apical micron-sized clathrin-coated endocytic tubules (ETs) that allow for the concentration and rapid internalization of soluble protein cargoes, while excluding resident apical membrane proteins. Dab2, an endocytic adaptor linking clathrin to scavenger receptors, controls the biogenesis and function of ETs by recruiting unconventional myosin Myo6 and by influencing dynamin activity. Our findings suggest that continuous pinching of vesicles from ETs allows for rapid, high-capacity internalization of dietary protein.

**Graphical abstract:** 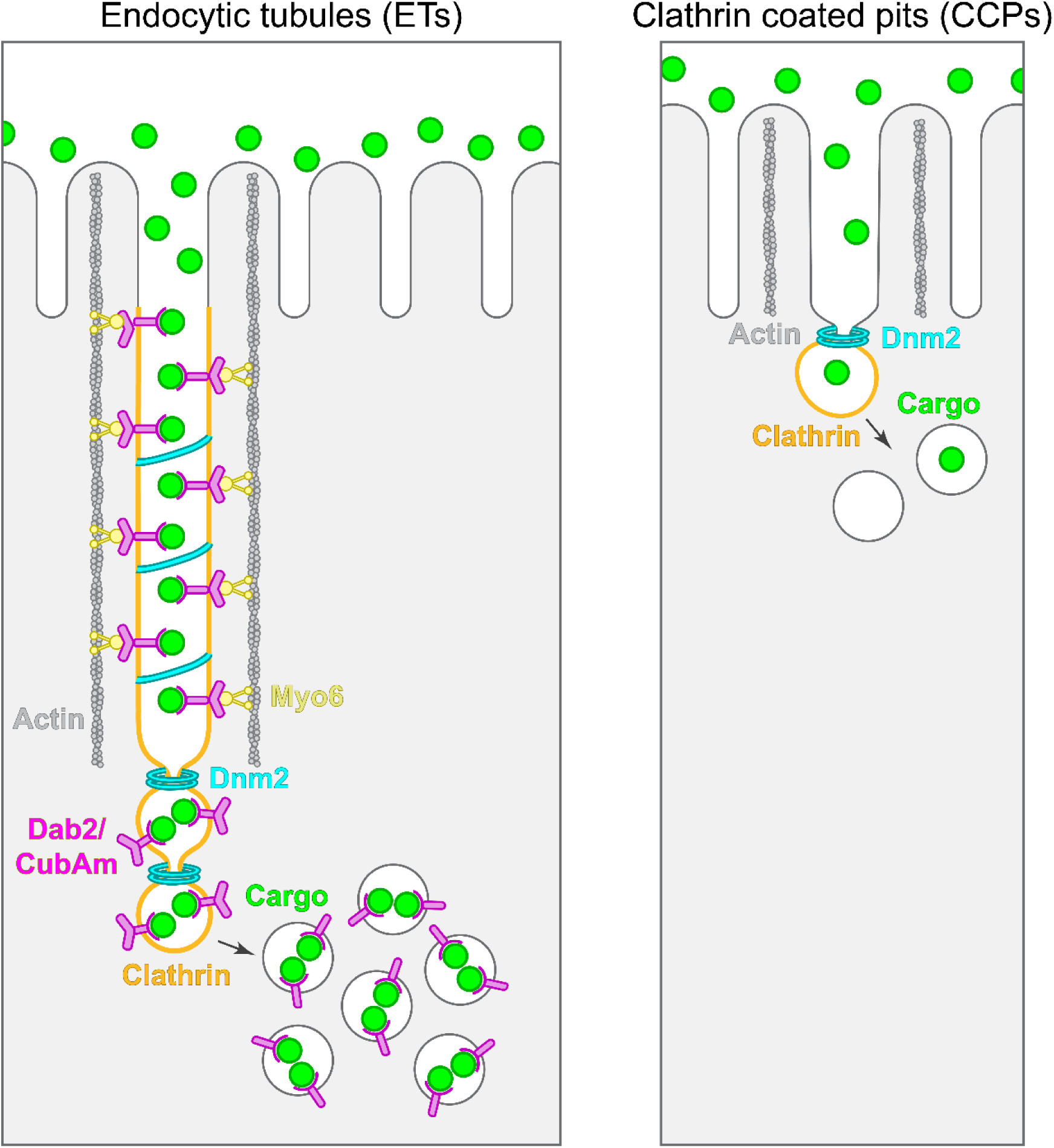

## Introduction

The intestine is the site of nutrient absorption in animals. In the mature intestine of vertebrates, the enterocyte apical membrane, known as the brush border, provides a highly specialized and expanded surface for nutrient uptake. Macronutrients are largely digested in the acidic lumen of the gastrointestinal track, and brush border localized transporters then facilitate the uptake of amino acids, sugars, and fatty acids into enterocytes (1, 2). The brush border itself also has important roles in digestion through the expression of membrane hydrolases that break down remaining macromolecules (1, 2), and brush border microvilli tips also continuously shed vesicles that harbor digestive enzymes (3). By contrast, in the immature intestine of neonates and stomachless fish such as zebrafish luminal proteolysis is highly inefficient due to relative high pH and low protease abundance (4–6). In mammals, these conditions allow for the preservation of antigens and antibodies present in milk (7) but also pose the challenge of absorption as proteins are essential nutrients.

Nutrient absorption in the neonatal mammalian gut relies on intracellular digestion by a population of enterocytes known as vacuolated or neonatal enterocytes (8–12). These cells are conserved between zebrafish and mammals and are specialized in the uptake and digestion of dietary proteins (13, 14). They are found in the ileum in mammals and in a homologous region of the posterior mid-gut in zebrafish (13–15). As these cells possess a prominent lysosomal vacuole and are present in non-mammalian vertebrates such as fish, we refer to them as Lysosome Rich Enterocytes (LREs).

Recent work has shown that dietary proteins are internalized in their native conformation by endocytosis into LREs, which then degrade proteins intracellularly in their large lysosomal vacuole (13). A fraction of internalized proteins undergo transcytosis via LREs and can be found in circulation. In mice, this process occurs largely in the jejunum via the same machinery found in LREs at the ileum and allows preservation of bioactive macromolecules to support neonatal development (14). In mammals, LREs are replaced by mature enterocytes at weaning (16–19) but in stomachless fishes like zebrafish, LREs remain in the intestine throughout adulthood and are essential for organismal growth and survival (13).

LREs internalize proteins by a form of receptor-mediated endocytosis that utilizes a broad-spectrum scavenger receptor complex consisting of cubilin (Cubn), its transmembrane linker amnionless (Amn), and the endocytic adaptor disabled 2 (Dab2) (13). While receptor-mediated endocytosis is effective at capturing diluted ligands, it is also subject to saturation when ligands are abundant and exceed the binding capacity of available receptors. Intriguingly, because Dab2 is a clathrin adaptor, it is unclear how clathrin mediated endocytosis (CME) could possibly fulfil the high demands of LRE endocytosis. Furthermore, while LREs are a small fraction of enterocytes (14, 20), they are solely responsible for dietary protein absorption, making the conundrum of how CME could sustain high-capacity protein uptake needed for organismal growth even more puzzling (11, 12, 21–25).

CME has been known for decades to be mediated by the iconic soccer ball like clathrin coated pit (CCP) structures. Internalization via CME occurs constitutively in most, if not all, cells to facilitate membrane recycling for normal plasma membrane maintenance and to allow for specialized processes such as antigen/antibody internalization and cell migration. CME occurs as a gradual process in which dozens of proteins are recruited to the plasma membrane over a period of 1-2 minutes, forming CCPs that ultimately undergo scission to be released from the plasma membrane into the cytoplasm (26). CME is initiated when clathrin subunits and adaptors cluster with cargoes at the plasma membrane, with recruitment of other proteins occurring sequentially thereafter (27–31). At synapses, where CME occurs rapidly, the hurdle of this rate-limiting initiation step is overcome by clathrin priming or pre-positioning (32). Although rapid CME has been extensively studied in neuronal synapses (33–36) little is known about the mechanisms mediating high-capacity CME in professional absorptive cells *in vivo*.

Here, using live microscopy kinetic assays in zebrafish, novel endogenous reporters for endocytic components, transmission electron microscopy (TEM), and genetic perturbations, we find that LREs utilize a unique form of CME that occurs through long, > 1 µm, endocytic tubules (ETs). ETs are continuous with the apical membrane, exposed to the intestinal lumen and serve to trap and concentrate protein cargoes. Remarkably, ETs are decorated with clathrin and adaptors along their length. ET biogenesis is regulated by Dab2 and the unconventional myosin Myo6. In the absence of Myo6, ETs are replaced by CCPs, but these are unable to sustain high-capacity uptake in LREs. Overall, our data reveal that ETs function as preformed, cargo-filled tubules that allow for continuous vesicle shedding at their tips, thus allowing LREs to internalize cargoes rapidly and with high-capacity via CME.

## Results

### LREs have a high capacity for selective endocytosis

Previous work has shown that LREs readily internalize luminal proteins and digest them intracellularly to support nutrient utilization and organismal growth in zebrafish and neonatal mice and identified a scavenger receptor, and select endocytic components needed for protein uptake (13). However, mechanistic understanding of the LRE endocytic processes and how these support the high internalization capacity underlying LRE function was lacking. To investigate these processes, we developed a microscopy-based kinetic assay to monitor LRE endocytosis in zebrafish larvae with high precision. To this end, we gavaged (Fig. 1A) 6 days post fertilization (dpf) larvae with GFP as protein cargo and FM4-64 (a lipophilic dye that incorporates into the plasma membrane). We then allowed endocytosis to proceed for 0.5-8 minutes before gavaging larvae with 4% PFA (Fig. 1B) and performed confocal imaging on intestinal cross-sections. Strikingly, GFP internalization into LREs was extremely rapid, becoming enriched intracellularly within only 0.5 minutes and reaching saturation within 3 min. (Fig. 1C, D, G, H). This high rate of endocytosis was accompanied by rapid plasma membrane turnover as FM4-64-labeled lipids were also strongly enriched intracellularly in LREs after only 0.5 minutes. (Fig. 1C, D, G, H). As a comparison, we performed the same analyses for enterocytes of the anterior midgut (AECs) and could not detect any protein endocytosis within the same time frame. Furthermore, the apical plasma membrane showed virtually no turnover (Fig. 1E-H). Together, these data show that LREs exhibit a high rate of apical endocytosis.

**Figure 1.**
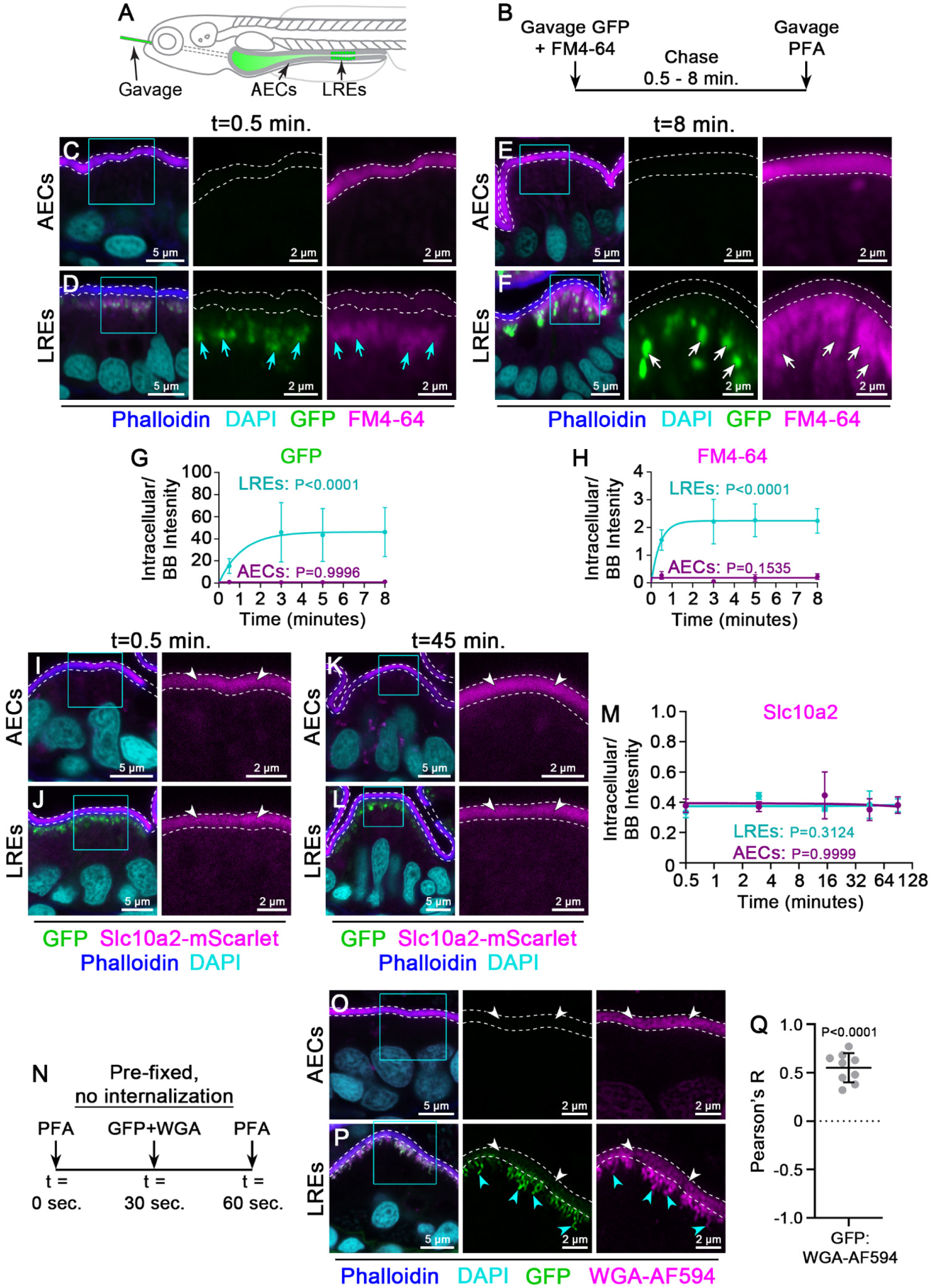
LREs have a high capacity for rapid and selective endocytosis. **(A)** Cartoon depicting gavage approach for studying luminal cargo uptake into zebrafish lysosome rich enterocytes (LREs) relative to anterior enterocyte controls (AECs). **(B)** Experimental design. **(C-F)** Confocal images of AECs (C, E) and LREs (D, F) of 6 dpf WT larvae gavaged with GFP (Green) and FM4-64 (Magenta), fixed at 0.5 minutes (C, D) and 8 minutes (E, F) post gavage and stained with Phalloidin-AF647 (F-actin, Blue) and DAPI (Cyan). Apical brush border (BB) is marked by dashed lines using the Phalloidin signal. Cyan arrows show internalized GFP and FM4-64 at 0.5 minutes (D), white arrows show internalized GFP and FM4-64 at 8 minutes (F). Cyan boxes are the regions of interest (ROIs) for magnified insets. **(G, H)** Quantification of intracellular / BB intensity of GFP (G) and FM4-64 (H) in LREs and AECs plotted against time (minutes). n ≥ 25 cells from ≥ 3 larvae. P values represent comparisons of t = 0.5 minutes versus t = 3 minutes and were calculated using Two-way ANOVA. Error bars are standard deviation (SD). **(I-L)** Confocal images of AECs (I, K) and LREs (J, L) of 6 dpf *TgKI(slc102a-mScarlet)^pd1308^* larvae gavaged with GFP (Green), fixed at 0.5 minutes (I, J) and 45 minutes (K, L) following GFP gavage and stained with phalloidin-AF647 (Blue) and DAPI (Cyan). BB is marked by dashed lines using the phalloidin signal. White arrowheads show Slc10a2-mScarlet localization at the BB. Cyan boxes are the ROIs for magnified insets. **(M)** Quantification of Intracellular / BB intensity of Slc10a2-mScarlet in LREs and AECs plotted against time (minutes). n ≥ 2 larvae per timepoint. P values represent comparisons of t = 0.5 minutes versus t = 3 minutes and were calculated using Two-way ANOVA. Error bars are SD. **(N)** Experimental design schematic. Brief pre-fixation of enterocytes is done with 4% PFA gavage for 30 seconds before GFP + WGA-AF594 gavage. **(O, P)** Confocal images of AECs (O) and LREs (P) of 6 dpf WT larvae gavaged with 4% PFA followed by GFP (Green) + WGA-AF594 (Magenta) gavage and then fixed and stained with phalloidin-AF647 (Blue) and DAPI (Cyan). BB is marked by dashed lines using the phalloidin signal. White arrowheads point to BB localization (O, P) and cyan arrowheads point to sub-apical tubular endocytic compartments labeled with GFP and WGA-AF594 (P). Cyan boxes are the ROIs for magnified insets. **(Q)** Quantification of Pearson’s R value for co-localization of luminally delivered protein cargo (GFP) and apical plasma membrane and derived compartments of LREs (WGA-AF594). n = 9 ROIs from 2 larvae. Data points are individual ROIs and P value was calculated using a one-sample t test. Error bars are SD.

Cargo internalization in LREs has been thought to be mediated by bulk endocytosis, which is non-selective and involves the engulfment of large plasma membrane domains (11, 21). Therefore, we wanted to test if LRE endocytosis is selective for soluble cargoes or if it causes internalization of resident apical membrane proteins. Previous work identified the apical bile acid transporter solute carrier family 10 member 2 (Slc10a2) to be expressed in ileocytes and anterior LREs (37, 38), making it an ideal candidate to examine in LREs alongside AEC control cells. We therefore generated an endogenously tagged allele for *slc10a2 (TgKI(slc10a2-mScarlet)^pd1308^)* and observed robust expression in ileocytes and anterior LREs with specific localization to their apical plasma membrane. To test whether Slc10a2 is internalized along with protein cargo, we gavaged 6 dpf *TgKI(slc10a2-mScarlet)^pd1308^* larvae with GFP and monitored its internalization over time. In contrast to the rapid internalization of GFP (Fig. 1G), Slc10a2-mScarlet remained restricted to the apical membrane in LREs (Fig. 1J, L, M) and control AECs (Fig 1I, K, M). Similarly, we also tested internalization of p75-GFP (an O-glycosylated apical membrane protein) (39) relative to mCherry and observed virtually no internalization of p75-GFP in LREs or AECs (Fig. S1 A-F, H). These results suggest that despite the high turnover of the apical plasma membrane (∼80% within less than 3 minutes) (Fig. 1H), resident apical membrane proteins are selectively excluded from LRE endocytosis.

Although we observed GFP cargo accumulating in intracellular vesicles within only 0.5 minutes after gavage (Fig 1D, G), it is unclear how it concentrates so rapidly in endosomes and what intermediate compartments are involved. To address this question, we modified our experimental design to briefly pre-fix the gut using a 0.5-minute gavage with 4% PFA to achieve complete inhibition of endocytosis, followed by cargo gavage (Fig. 1N). Using this experimental design, we gavaged 6 dpf WT zebrafish larvae with GFP and WGA-AF594 (a fluorescently conjugated lectin that labels glycosylated proteins abundant in the plasma membrane). Surprisingly, we found that when endocytosis is blocked, GFP is present in tubular compartments below the apical membrane of LREs (Fig 1P). By contrast, no such compartments were observed in AECs (Fig. 1O). Moreover, because WGA overlaps with GFP in these tubular compartments, these data suggest that luminally delivered protein cargo (GFP) is lodged within long invaginating structures derived from the apical membrane (Fig. 1Q).

### LREs harbor endocytic tubules (ET) that transport luminal cargos to early endosomes

Our data show that in LREs endocytic cargo traverses a tubular compartment before internalization. To determine whether these structures are truly continuous with the apical plasma membrane, we exploited the pH difference between the gut lumen and intracellular vesicles (40). We generated a pHluorin-TagRFPt pH sensor fusion protein, which we calibrated by generating a standard curve of pH-dependent intensity profiles (Fig. 2A, B). We found that pHluorin exhibits maximal intensity at pH 8, which corresponds to gut luminal pH in zebrafish larvae and is quenched at pH 6, which corresponds to early endosome pH (5, 41). By contrast, tagRFPt exhibits maximal intensity at pH 6-6.5 and reduced fluorescence at pH 8 (Fig. 2A, B), indicating that our sensor can be used to determine whether apical tubular compartments are luminally exposed. Next, we gavaged our pH sensor into 6 dpf WT larvae and performed live imaging 10-20 minutes after gavage. We observed a 13.6-fold higher pHluorin/TagRFPt intensity ratio in apical tubular structures compared to intracellular endosomes, suggesting that the tubular structures responsible for initial concentration of protein cargo have a neutral pH like the gut lumen. By contrast, pHluorin fluorescence was quenched in endosomes as it encountered an acidic pH (Fig 2C, D). Together, our results demonstrate that luminally delivered protein cargo is first concentrated in tubular invaginations that are continuous with the apical plasma membrane of LREs. Henceforth, we refer to these unique structures as endocytic tubules (ETs).

**Figure 2.**
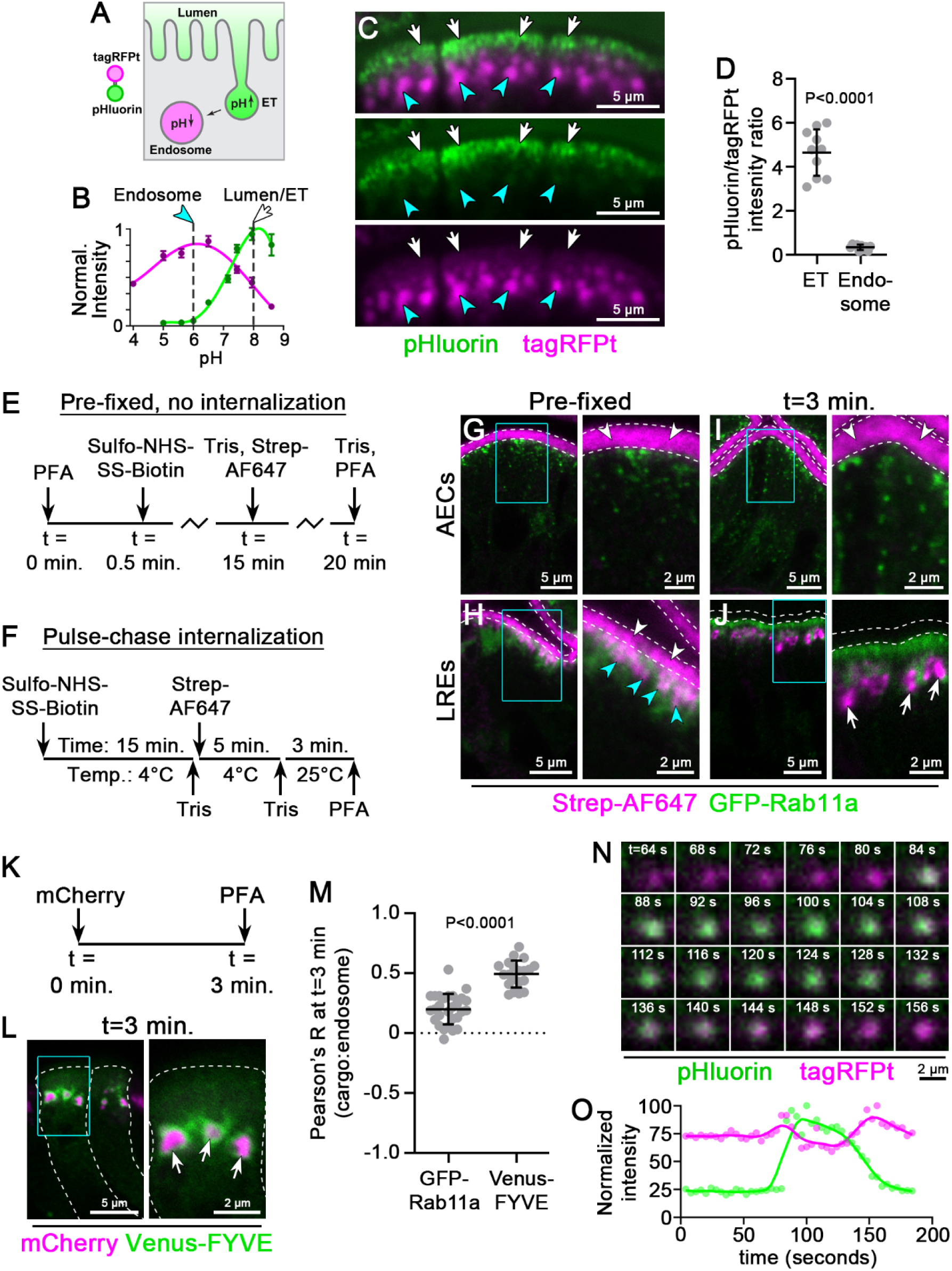
LREs harbor endocytic tubules (ET) that transport luminal cargos to early endosomes. **(A)** Cartoon depicting pH dependent fluorescence of the pHluorin-tagRFPt sensor. **(B)** Standard curve showing normalized fluorescence intensity of pHluorin-tagRFPt vs pH. Data analysis is described in the Materials and Methods section. Error bars are standard error of the mean (SEM). Green arrowhead arrow indicates endosomal pH (∼6) and white arrow luminal pH (∼8). **(C)** Live confocal image of apical region of 6 dpf WT LREs showing high pHluorin signal in luminally exposed endocytic tubules (ETs, white arrows) compared to intracellular regions with high tagRFPt signal in endosomes (cyan arrowheads). **(D)** Quantification of pHluorin/tagRFPt fluorescence intensity ratios in ETs and endosomes. n = 10 cells from 2 larvae. P value was calculated using Welch’s t test. Error bars are SD. **(E, F)** Experimental design. **(G-J)** Confocal images of AECs (G, I) and LREs (H, J) of 6 dpf WT *TgKI(GFP-rab11a)^pd1124^*larvae gavaged as described in panel E (G, H) and panel F (I, J). BB is marked by dashed lines. White arrowheads point to BB localization (G, H, I), cyan arrowheads point to sub-apical ETs marked by Streptavidin-AF647 signal beneath the BB (H). White arrows point to Streptavidin-AF647 signal in endosomes after a 3 minute chase period (J). Cyan boxes are the ROIs for magnified insets. **(K)** Experimental design to study gavaged luminal cargo (mCherry) localization with respect to endosomes (marked by Venus-FYVE expression in 6 dpf *Tg(hsp:Venus-FYVE)^pd1046^* larvae) after a 3-minute chase. **(L)** Confocal images of LREs from 6 dpf *Tg(hsp:Venus-FYVE)^pd1046^* larvae gavaged with mCherry and fixed with 4% PFA gavage after a 3-minute chase. Cyan box is the ROI for the magnified inset. **(M)** Quantification of co-localization of apical membrane (marked by Streptavidin-AF647) with recycling endosomes (shown in panel J) and luminal cargo (mCherry) with Venus-FYVE marked early endosomes (shown in panel L) after a 3-minute chase. n ≥ 20 cells from ≥ 4 larvae. P value was calculated using an unpaired t test. Error bars are SD. **(N)** Time-lapse live imaging of an LRE early endosome after gavage with pHluorin-tagRFPt, showing pH dependent quenching of pHluorin after a fusion event. **(O)** Normalized intensity values of pHluorin and tagRFPt fluorescence from the time-lapse shown in panel N

Our data so far show that LREs use ETs to rapidly internalize cargo, which then seems to accumulate in endosomes within seconds after gavage. To investigate the fate of internalized cargo, we performed pulse-chase assays using a cell surface biotinylation approach. We gavaged Sulfo-NHS-SS-Biotin to label apical membrane proteins, quenched the reaction with excess Tris, and then gavaged Streptavidin-AF647 to visualize labeled proteins. As before, when pre-fixed with a brief PFA gavage (Fig. 2E), only the apical membrane was labeled with this scheme in control AECs (Fig. 2G), whereas ETs were also strongly labeled in LREs (Fig. 2H). Next, for pulse-chase analysis, we performed the same labeling scheme in live larvae but allowed labeled proteins to be internalized for 3 minutes before processing (Fig. 2F). Control AECs showed no internalization of labeled apical membrane proteins (Fig. 2I). By contrast, we observed high turnover of the apical membrane in LREs, with labeled proteins accumulating intracellularly in a compartment distinct from recycling endosomes, marked with GFP-Rab11a (Fig. 2J). We next performed similar endocytosis assays in larvae expressing Venus-FYVE, an early endosome marker (42, 43), and observed robust cargo accumulation within the compartment 3 minutes after gavage (Fig. 2K-M). Furthermore, live time-lapse imaging of larvae gavaged with our pH sensor showed that cargo transported from ETs is delivered to acidic early endosomes, where pHluorin is quenched within 1 minute of fusion (Fig. 2N, O). Together, these results demonstrate that LREs possess ETs open to the gut lumen that rapidly deliver cargo to early endosomes.

### ETs are Clathrin-coated plasma membrane invaginations that internalize luminal proteins in LREs

To investigate the morphology and ultrastructure of ETs, we performed Transmission Electron Microscopy (TEM) analysis of LREs compared to AECs from 6 dpf WT larvae. Whereas in AECs endocytic pits were sparse and rare (Fig. 3A-D), LREs showed extensive tubular invagination of the apical membrane originating from the intermicrovillar cleft (Fig. 3E-L). These structures seem to extend to depths hundreds of nanometers below the apical membrane (Fig. 3M). Strikingly, these invaginations were continuously lined by a spiked or shadowed appearance (Fig. 3F-H, J-L, arrowheads), which is indicative of clathrin coated membranes (44, 45). These clathrin-like features seem to decorate ETs along their entire length (Fig. 3J-L, arrowheads), with vesicles appearing to pinch from mostly from the base but can also be found at other sites along the length ETs (Fig. 3F-H). Interestingly, we found that LREs possess shorter microvilli compared to AECs (Fig. 2O), suggesting that ETs are formed at the expense of microvilli. Together, these data reveal that ETs are long clathrin-coated invaginations that are continuous with the apical membrane.

**Figure 3.**
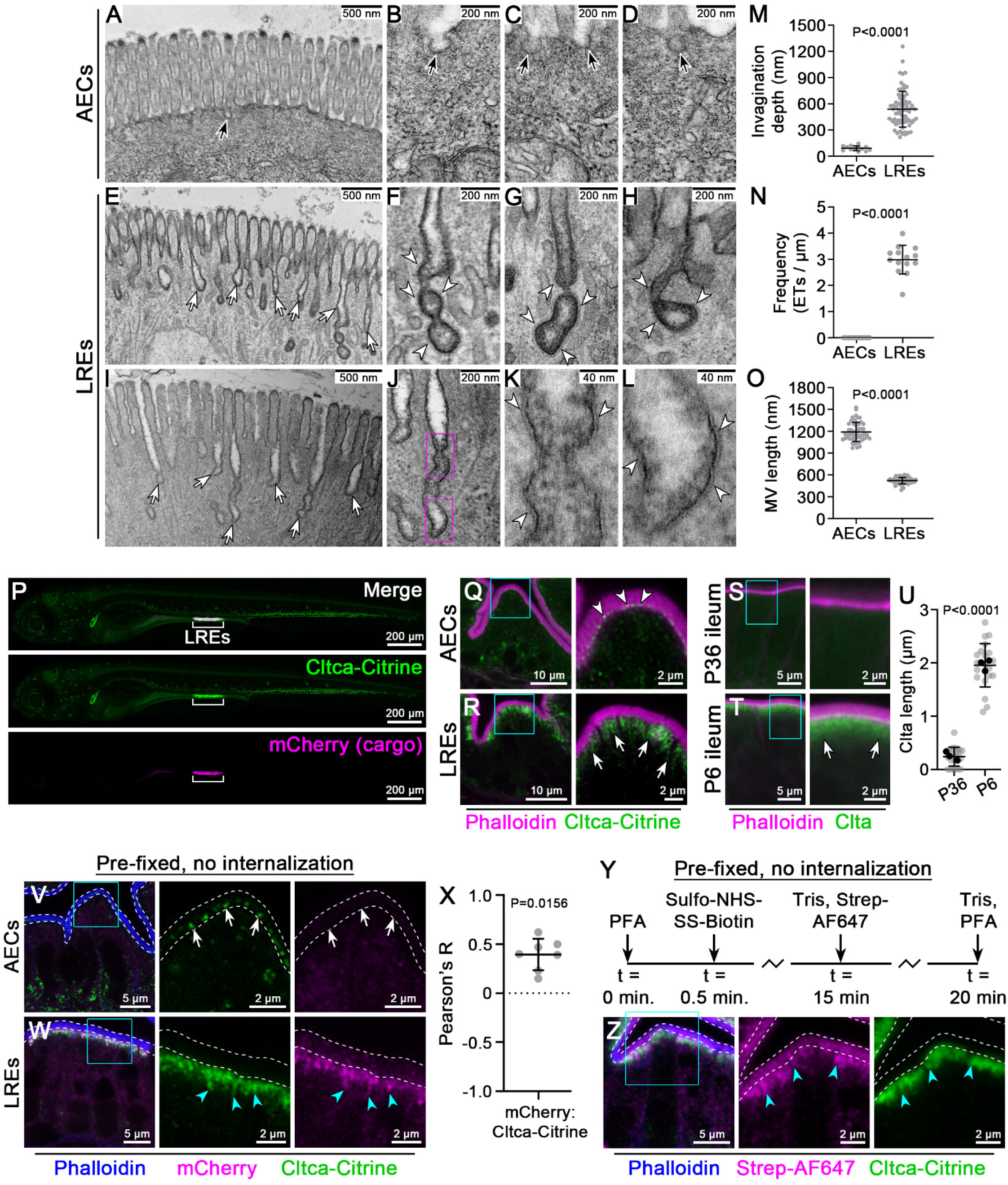
ETs are long Clathrin-coated plasma membrane invaginations exposed to the intestinal lumen. **(A-L)** Transmission electron microscopy (TEM) images of 6 dpf WT AECs (A-D) and LREs (E-L). Black arrows show clathrin coated pits (CCPs) in AECs (A-D), white arrows point to ETs in LREs (E, I). White arrowheads point to electron dense patterns along ETs, denoting clathrin-coats (F-H, K, L). Magenta boxes are the ROIs for magnified insets in panels K and L. **(M)** Endocytic invagination depth measurements from TEM images of AECs (n = 2 larvae) and LREs (n = 3 larvae). Error bars are SD. **(N)** Number of ETs per micrometer of apical surface measured from TEM images of AECs (n = 2 larvae) and LREs (n= 2 larvae). Error bars are SD. **(O)** Microvilli length measurements from TEM images of AECs (n = 3 larvae) and LREs (n = 5 larvae). Error bars are SD. P values for panels M-O were calculated using Mann-Whitney tests. **(P)** Maximum intensity projections of a 6 dpf *Gt(cltca-citrine)^ct116a^* larva 15 minutes after gavage with mCherry. Brackets denote LREs. **(Q, R)** Deconvolved confocal images of AECs (Q) and LREs (R) in 6 dpf *Gt(cltca-citrine)^ct116^* larvae stained with Phalloidin-AF594 (Magenta). White arrowheads show CCPs in AECs (Q), whereas white arrows show ETs in LREs (R) that are marked by Cltca-citrine (green). Cyan boxes are the ROIs for magnified insets. **(S, T)** Deconvolved confocal images of P6 and P36 WT mice cryosections of the ileal region stained with Clta (Clathrin light chain) Antibody (Green) and Phalloidin-AF647 (Magenta). White arrows show ETs in P6 ileum that harbors LREs (T). Cyan boxes are the ROIs for magnified insets. **(U)** Quantification of the length of Clta antibody signal beneath the BB (marked by Phalloidin-AF647) in the ileum at P36 (post-weaning, n = 3 mice) and P6 (pre-weaning, n = 3 mice). Gray data points are cells pooled from multiple animals, while black data points are mean lengths from individual animals. Error bars are SD. P value was calculated using a nested t test. **(V, W)** Deconvolved confocal images of AECs (V) and LREs (W) of 6 dpf *Gt(cltca-citrine)^ct116a^* larvae gavaged with 4% PFA to block endocytosis followed by mCherry gavage (Magenta), post-fixation, and staining with Phalloidin-AF647 (Blue). BB is marked by dashed lines using the Phalloidin-AF647 signal. White arrows point to Cltca-citrine labeled CCPs in AECs (V) and cyan arrowheads point to Clta-citrine labeled ETs (W). Cyan boxes are the ROIs for magnified insets. **(X)** Quantitation of Pearson’s R value for colocalization of luminally delivered protein cargo (mCherry) and Cltca-citrine labeled ETs in *Gt(cltca-citrine)^ct116a^*larvae. n = 7 ROIs from 4 larvae. Data points are individual ROIs. Error bars are SD. P value was calculated using a one-sample Wilcoxon test. **(Y)** Experimental design. **(Z)** Confocal images of LREs from 6 dpf *Gt(cltca-citrine)^ct116a^*larvae gavaged with 4% PFA to block endocytosis followed by cell surface biotinylation labeling (Streptavidin-AF647, Magenta), post-fixation, and staining with Phalloidin-AF594 (Blue). BB is marked by dashed lines using the Phalloidin signal. Cyan arrowheads point to Cltca-citrine labeled ETs positive for the apical membrane (Streptavidin-AF647). Cyan boxes are the ROIs for magnified insets.

To further investigate ETs, we performed live imaging of whole larvae expressing endogenously tagged clathrin heavy chain (Cltca-Citrine, which is expressed in the Tg*Gt(cltca-citrine)^ct116a^* line) (46). While Cltca-Citrine is expressed in multiple organs, it is most highly enriched in LREs, where it overlapped with mCherry after gavage (Fig. 3P, brackets). High resolution imaging on intestinal cross-sections revealed that Cltca-Citrine strongly labeled ETs in LREs (Fig. 3R), consistent with our TEM analysis (Fig. 3E-L). By contrast, AECs showed sparse punctate labeling interspersed along the base of the apical membrane, likely representing clathrin-coated pits (CCPs) (Fig. 3Q). We previously found that LREs are present in the ileum of neonatal mice at suckling stages and are then lost post-weaning (14). To determine whether clathrin-coated ETs are also present in mouse LREs, we compared clathrin light chain (Clta) immunolocalization in ileal enterocytes in pre-weaning stage (P6) and post weaning stages (P36). We found that P6 LREs exhibited higher levels and distinct localization of apical Clta compared to P36 ileocytes (Fig. 3S, T), with LREs showing a clathrin rich region extending hundreds of nm below the apical membrane (Fig. 3U). This is consistent with early TEM data showing apical invaginations in neonatal mice and rat enterocytes (11, 12, 21–23). These results suggest that clathrin-coated ETs are conserved in mammalian LREs.

We next tested whether the clathrin-labeled structures observed in LREs are in fact the ETs that transport endocytic cargo. To this end, we gavaged mCherry in 6 dpf *TgGt(cltca-citrine)^ct116a^* larvae that were briefly pre-fixed by a 0.5-minute PFA gavage to prevent cargo internalization. Similar to GFP in pre-fixed conditions (Fig. 1Q), mCherry cargo was highly enriched in apical ETs labeled by Cltca-citrine (Fig. 3V-X). Further, we performed cell surface biotinylation by gavage and observed that Streptavidin-AF647 labeled ETs that were marked by Cltca-Citrine (Fig. 3Y, Z). Together, our results show that ETs are long clathrin-coated invaginations that mediate the rapid uptake of luminal cargo and select apical membrane proteins.

### Internalization through ETs is Dynamin- and Clathrin-dependent

Our data suggest that LREs use a unique form of CME that occurs through ETs. To further investigate this mechanism, we next functionally tested the roles of key endocytic components in this process. Dab2, a clathrin adaptor (47), is critical for protein and fluid phase endocytosis in LREs (13). To determine whether Dab2 associates with Clathrin on ETs, we generated an endogenously tagged allele, *TgKI(dab2-mScarlet)^pd1292^*. We found that the spatial expression patterns of Dab2-mScarlet in whole larvae closely matched that of Cltca-Citrine, with highest enrichment seen in LREs and the pronephros (Fig. 4A, B). We also generated an endogenously tagged allele for dynamin2a, *TgKI(GFP-dnm2a)^pd1274^*, and found that it was similarly enriched in LREs (Fig. 4C). Next, we performed high resolution imaging on intestinal cross sections and found that Dab2-mScarlet strongly overlapped Cltca-citrine on ETs (Fig. 4 D). We also observed similar enrichment of GFP-Dnm2a subapically at ETs, which could also be visualized with Phalloidin labeling of F-actin (Fig. 4E).

**Figure 4.**
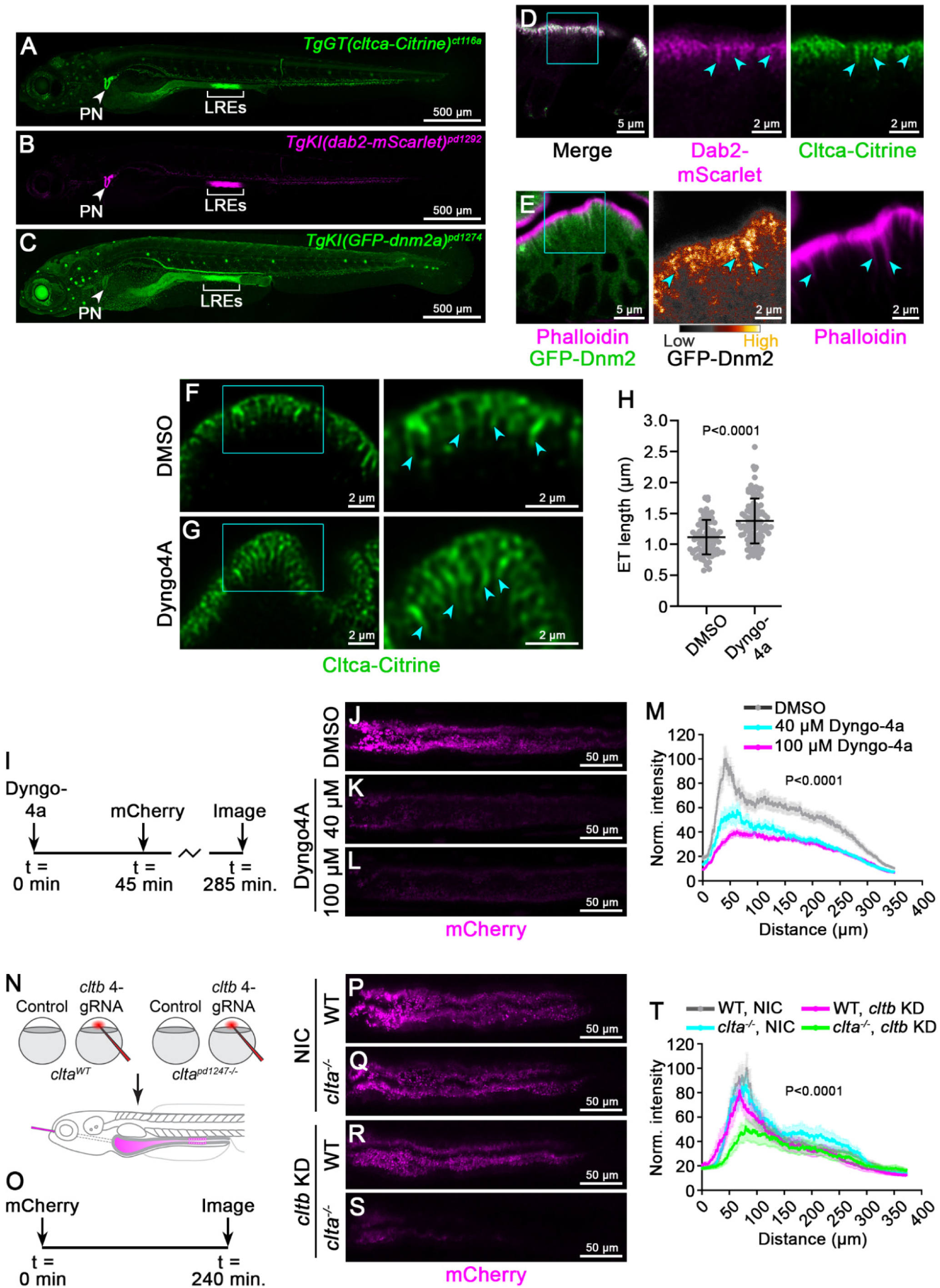
Internalization through ETs is Clathrin- and Dynamin-dependent. **(A-C)** Maximum intensity projections of 6 dpf *Gt(cltca-citrine)^ct116a^* (A), *TgKI(dab2-mScarlet) ^pd1292^* (B), and *TgKI(GFP-dnm2a)^pd1274^* (C) larvae showing high expression in LREs (brackets) and pronephros (arrowheads). **(D)** Deconvolved confocal image of 6 dpf *Gt(cltca-citrine)^ct116^*; *TgKI(dab2-mScarlet) ^pd1292^* LREs. Cyan arrowheads point to Clta-citrine and Dab2-mScarlet labeled ETs. Cyan box is the ROI for magnified inset. **(E)** Confocal images of 5 dpf *TgKI(GFP-dnm2a)^pd1274^* LREs stained with Phalloidin-AF647 (magenta, right panel). Cyan arrowheads point to ETs. Middle panel of GFP-Dnm2a is pseudo-colored using the smart lookup table (LUT) in ImageJ. Cyan box is the ROI for magnified inset. **(F, G)** Deconvolved confocal images of 6 dpf *Gt(cltca-citrine)^ct116^*LREs after gavage with DMSO vehicle control (F) or Dyngo4a (Dynamin inhibitor) (G). Cyan arrowheads point to Cltca-citrine labeled ETs. Cyan boxes are the ROIs for magnified insets. **(H)** Quantification of ET length. n = 4 larvae (DMSO) or n = 6 larvae (Dyngo4a). Data points are individual ETs. P value was calculated using a Mann-Whitney test. **(I)** Experimental design. **(J-L)** Maximum intensity projections of confocal stacks of LRE regions showing mCherry uptake in 6 dpf WT larvae treated with DMSO vehicle control (J), 40 μM Dyngo4a (K), or 100 μM Dyngo4a. **(M)** Quantification of total mCherry uptake from panels J-L. n ≥ 17 larvae per group. Error bars are SEM. P value was calculated using Two-way ANOVA. **(N, O)** Schematic diagram showing experimental design for testing luminal protein cargo uptake upon loss of Clathrin function. Embryos collected from in-crosses of c*lta^pd1247+/−^* heterozygous parents were injected with a pool of 4 gRNAs targeting multiple regions of the *cltb* gene (clathrin light chain b, paralog of *clta*). At 6 dpf, these larvae were gavaged mCherry (O) and genotyped after imaging. **(P-S)** Maximum intensity projections of confocal stacks of LRE regions at 6 dpf showing mCherry uptake for the genotypes described in panel N. **(T)** Quantification of total mCherry uptake for panels P-S. n ≥ 13 larvae per genotype). Error bars are SEM. P value was calculated using Two-way ANOVA.

To functionally test if LRE endocytosis though ETs is dynamin-dependent, we inhibited dynamin activity using Dyngo4a, which blocks the GTP hydrolysis step required for vesicle scission (48). Consistent with previous reports showing tubular elongation of CCPs upon dynamin inhibition (27, 35), we observed that ETs in *TgGt(cltca-citrine)^ct116a^* larvae gavaged with Dyngo4a were significantly longer than those of controls (Fig. 4F-H). Next, we performed endocytosis assays by pre-treating larvae with Dyngo4a or DMSO as a control for 45 minutes and then gavaging with mCherry (Fig. 4I). We found that Dyngo4a treatment caused a dose-dependent inhibition of mCherry endocytosis (Fig. 4J-M), indicating that dynamin activity is critical for endocytosis in LREs.

To test if Clathrin present on ETs has a functional role in LRE endocytosis, we generated a clathrin light chain mutant by targeting the *clta* gene using CRISPR/Cas9 (Fig. S2A). This *clta* mutant allele, *clta ^pd1247^,* has a 10 bp insertion that results in a premature stop codon and causes reduced *clta* mRNA levels (Fig. S2C, D). We first gavaged *clta^pd1247^* mutants with mCherry to assay cargo internalization but did not observe any reductions compared to WT control siblings (Fig. 4P, Q). We reasoned that a lack of a cargo uptake phenotype in *clta^pd1247^* mutants may be due to the activity of its paralog, *cltb*. Therefore, we performed knockdown (KD) of *cltb* using a CRISPR gRNA pool targeting 4 exons of the *cltb* gene (Fig. S2E) in *clta^pd1247^* mutants and WT siblings (Fig. 4N, O). While *cltb* KD alone did not show any reduction in the mCherry uptake in LREs, *clta^pd1247^*mutants injected with the *cltb* CRISPR gRNA pool showed a strong reduction in mCherry cargo uptake compared to controls (Fig. 4R, S). Together, our results indicate that LRE endocytosis through ETs is dynamin- and clathrin-dependent.

### Myosin VI is required for ET biogenesis and endocytosis in LREs

To better understand how cargo uptake through ETs is regulated, we analyzed transcriptome datasets of LREs and the rest of the intestinal epithelium (IECs) we previously generated (13) to identify genes potentially related to endocytosis function and with enriched expression in LREs. Similar to *dab2, cubn,* and *amn,* which encode the endocytic scavenger receptor complex crucial for endocytosis in LREs (13), *myo6a* was highly upregulated (Log2FC = 2.23) in LREs (Fig. 5A). This was notable because *myo6a* encodes myosin VI (Myo6), which is a Dab2-interacting protein (49, 50) and an unconventional myosin that is a minus end directed actin motor protein (51). Interestingly, enterocytes exhibit highly polarized apical F-actin orientation, with plus ends anchored in microvilli tips and minus ends extending towards the cytoplasm (52–55). Imaging LREs, we observed long F-actin bundles extending beneath microvilli in LREs (Fig. 4D) and microfilaments adjacent to ETs in or TEM analysis (Fig. 3I). We therefore hypothesized that Myo6 might function with Dab2 to regulate LRE endocytosis and possibly ET biogenesis.

**Figure 5.**
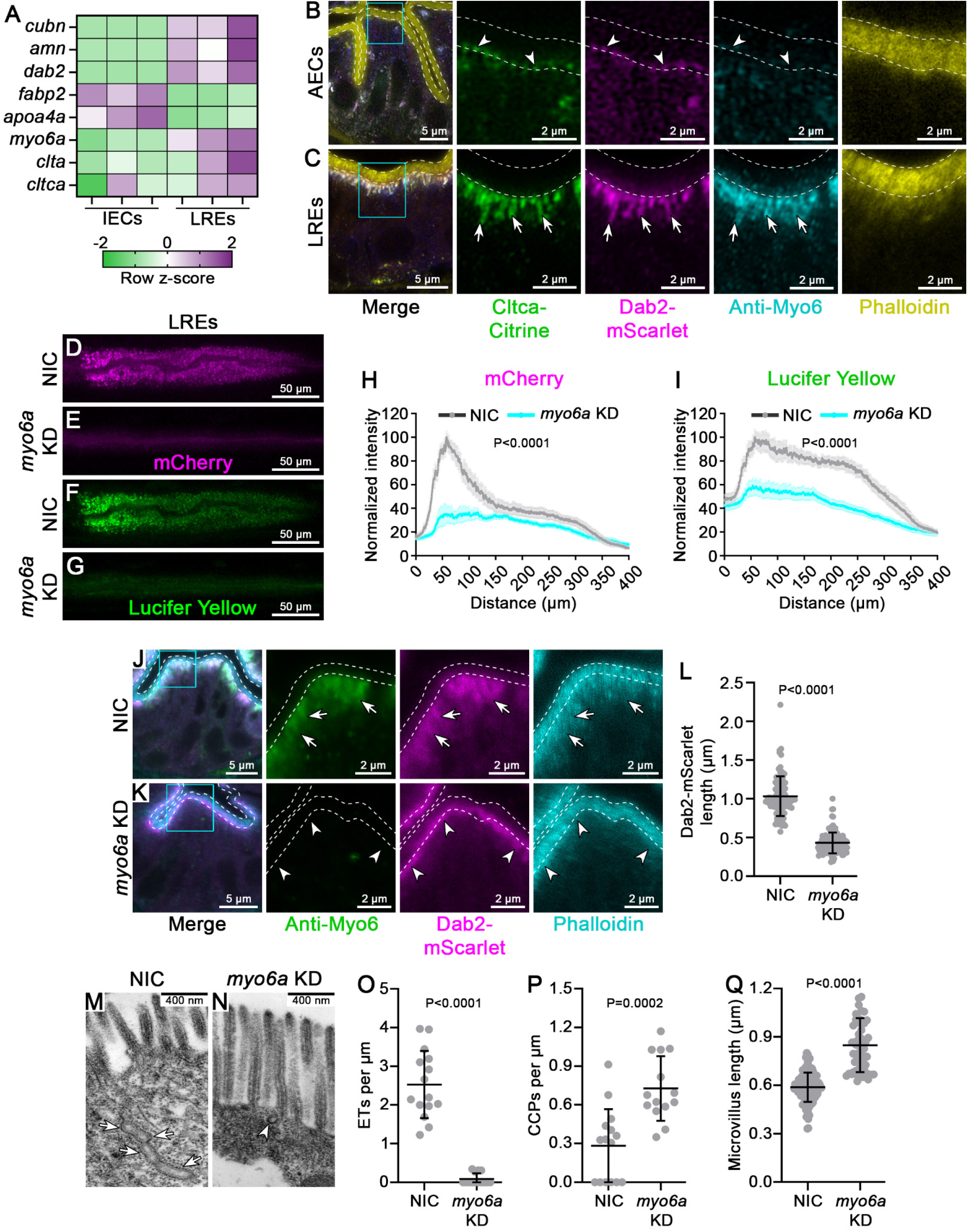
Myosin VI is required for ET biogenesis and endocytosis in LREs. **(A)** Heat map showing relative expression of selected genes in AECs (left) and LREs (right). Primary data was obtained from our published dataset (13). **(B, C)** Confocal images of AECs (B) and LREs (C) of 6 dpf *Gt(cltca-citrine)^ct116a^*; *TgKI(dab2-mScarlet)^pd1292^* larvae stained with Phalloidin-AF647 (yellow) and anti-Myo6 antibody (Cyan). Apical BB is marked by white dashed line using Phaloidin-AF647 staining as a reference. White arrowheads point to CCPs in AECs (B) and white arrows indicate co-localization of Myo6 and Dab2 with Cltca-citrine-labeled ETs. Cyan boxes are the ROIs for magnified insets. **(D-G)** Maximum intensity projections of confocal stacks showing mCherry (D, E) and Lucifer yellow (LY) (F, G) uptake in LRE regions of 6 dpf non-injected control (NIC) (D, F) and *myo6a* knockdown (KD) (E, G) larvae 4 hours after gavage. **(H, I)** Quantification of total mCherry (H) and LY (I) uptake for panels D and E. n ≥ 14 larvae. Error bars are SEM. P values were calculated using Two-way ANOVA. **(J, K**) Confocal images of LREs from NIC (J) and *myo6a* KD (K) 6 dpf *TgKI(dab2-mScarlet)^pd1292^*larvae stained with anti-Myo6 antibody (Green) and Phalloidin-AF647 (Cyan). White arrows show ETs decorated with Myo6 and Dab2-mScarlet in control LREs (J), and white arrowheads show loss of Myo6 Ab staining and a restricted Dab2-mScarlet localization in *myo6a* KD LREs (K). Cyan boxes are the ROIs for magnified insets. **(L)** Quantification of the length of Dab2-mScarlet on ETs from panels J and K. Data points are individual ETs from n ≥ 6 larvae. Error bars are SD. P values were calculated using a Mann-Whitney test. **(M, N)** TEM images of LREs from 6 dpf NIC control (M) and *myo6a* KD larvae (N). White arrows show a clathrin coated ET in NIC larvae (M), and white arrowhead shows a CCP in *myo6a* KD larvae. **(O)** Quantitation of ETs per micrometer of apical surface of LREs from TEM analysis. About 40 mm of apical surface analyzed for each group from multiple TEM images of NIC (n = 2 larvae) and *myo6a* KD (n = 3 larvae). Data points are values from individual images. **(P)** Quantification of CCPs per micrometer of apical surface of LREs in 6 dpf NIC and *myo6a* KD larvae as done as in panel O. n = 2 larvae (NIC) and n = 3 larvae (*myo6a* KD). Data points are values from individual images. **(Q)** Length of microvilli in LREs of 6 dpf NIC and *myo6a* KD larvae measured using TEM images. Data points are individual microvilli. For panels O-Q, error bars are SD, and P values were calculated using a Mann-Whitney test.

To investigate the role of Myo6 in LREs, we first performed *in situ* hybridization chain reaction (HCR) staining (56) to confirm that *myo6a* expression is indeed enriched in LREs (Fig. S3 A). Similar to endogenously tagged Dab2 and Cltca (Fig. 4A, B), we found robust expression of *myo6a* in LREs, as well as in the pronephros (Fig. S3A). To examine the subcellular localization of Myo6, we next performed antibody staining on cross sections of 6 dpf *TgGt(cltca-citrine)^ct116a^; TgKI(dab2-mScarlet)^pd1292^* larvae. We found that Myo6 strongly overlapped with Dab2-mScarlet on Cltca-citrine labeled ETs (Fig. 5C). Expression and localization of Myo6 was specific to LREs as control AECs, which do not form ETs, showed negligible Myo6 labeling (Fig. 5B, C), consistent with our HCR data (Fig. S3A) and RNAseq data (13).

Because Myo6 tightly associated with ETs and is a known Dab2-interacting protein (49, 50, 57–59), we next tested if Myo6a function is required for LRE endocytosis. To this end, we performed *myo6a* KD using a CRISPR gRNA pool that resulted in the loss of Myo6 antibody labeling compared to WT controls (Fig. 5J, K). Upon myo*6a* KD, we observed a sharp reduction in uptake of both mCherry protein cargo (Fig. 5D, E, H) and Lucifer Yellow fluid phase cargo (Fig. 5F, G, I) compared to WT control larvae, indicating that Myo6a is required for ET-mediated endocytosis. We next investigated if loss of Myo6 affects Dab2 localization. We found that upon *myo6a* KD, Dab2-mScarlet became spatially restricted to the base of microvilli and failed to label ETs (Fig. 5J, K, L). Subapical F-actin cables were also less organized in *myo6a* KD larvae (Fig. 5J, K). Overall levels of Dab2-mScarlet, however, appeared unchanged in *myo6a* KD larvae, suggesting that Myo6 is not required for the association of Dab2 with the apical membrane in LREs. These results suggest that Myo6 is required for ET biogenesis and function and that in their absence LRE endocytosis is strongly inhibited.

Next, to further investigate how Myo6 regulates ET biogenesis, we performed TEM analysis of LRE cross-sections from 6 dpf *myo6a* KD and WT control larvae. Strikingly, we found that ET formation was strongly reduced in *myo6a* KD larvae, where ET density dropped from about 2.5 per µm in WT controls to almost none in *myo6a* KD larvae (Fig. 5M, N, O). Interestingly, impaired ET formation in *myo6a* KD larvae was accompanied by an increase in CCP density (Fig. 5M, N, P). We also observed significantly longer microvilli in LREs upon *myo6a* KD compared control larvae (Fig. 5M, N, Q), consistent with the notion that ETs form at the expense of microvilli (Fig. 3O, Fig. S2A-F). Together, our results show that ET biogenesis is Myo6-dependent and suggest that, although endocytic components still assemble at the apical membrane as CCPs after loss of Myo6, CCPs cannot sustain the high rate of endocytosis characteristic of LREs.

### Dab2 regulates Myosin6 and Dynamin localization to ETs and is required for ET morphology

Because Dab2-mediated targeting of Myo6 to clathrin coated structures has been shown to facilitate endocytosis (58, 59), we next investigated how Dab2 influences Myo6 localization in LREs using CRISPR gRNA pool KD of *dab2*. Consistent with the *dab2* mutant phenotype (13), *dab2* KD strongly reduced mCherry endocytosis in LREs after gavage of 6 dpf larvae (Fig. S4E, F). Next, we performed *dab2* KD on *TgKI(dab2-mScarlet)^pd1292^; TgGt(cltca-citrine)^ct116a^* larvae to analyze Myo6 localization with antibody staining at 6 dpf. We found that loss of Dab2 (confirmed by loss of Dab2-mScarlet signal) partially uncoupled Myo6 from ETs (Fig. 6A-C). Interestingly, we also observed a significant reduction in ET density upon *dab2* KD, with remaining ETs showing increased length (Fig. 6A, B, D, E).

**Figure 6.**
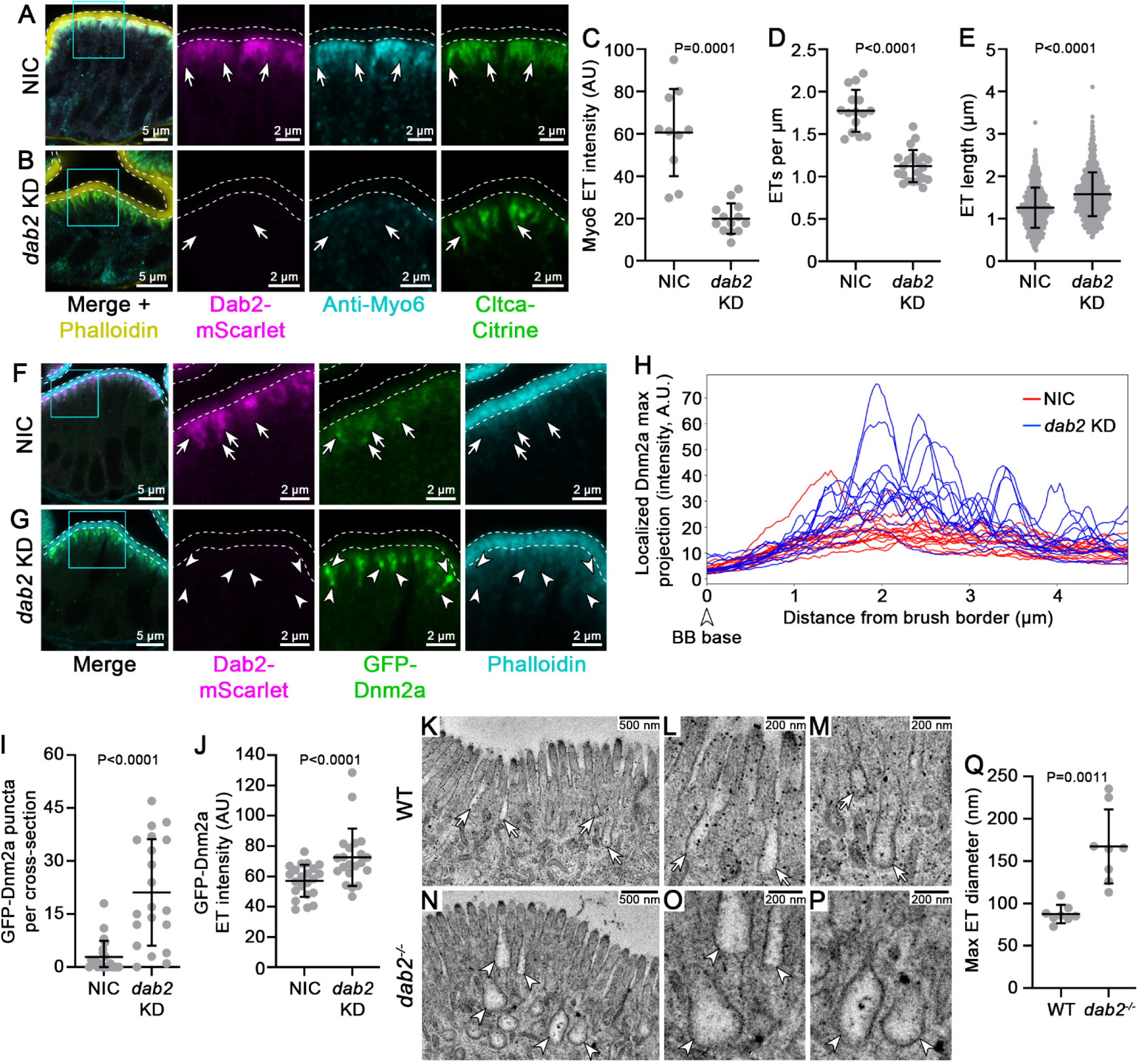
Dab2 regulates ET morphology and Myo6a recruitment to ETs. **(A, B)** Confocal images of 6 dpf *Gt(cltca-citrine)^ct116a^*; *TgKI(dab2-mScarlet)^pd1292^* NIC (A) and *dab2* KD (B) LREs stained with anti-Myo6 antibody (Cyan) and Phalloidin-AF647 (yellow), showing co-localization of Myo6 (Cyan) and Dab2-mScarlet (Magenta) with Cltca-citrine-labeled ET (Green) in control (A) but not in *dab2* KD (B). White arrows point to ETs. Cyan boxes are ROIs for magnified insets. **(C)** Quantitation of fluorescence intensity of Myo6 on ETs in NIC and *dab2* KD 6 dpf *Gt(cltca-citrine)^ct116a^*; *TgKI(dab2-mScarlet)^pd1292^* larvae. Data points are means of all ROIs for individual larvae. n ≥ 10 larvae per group. **(D)** Quantitation of number of ETs per micrometer of apical surface in NIC and *dab2* KD 6 dpf *Gt(cltca-citrine)^ct116a^*; *TgKI(dab2-mScarlet)^pd1292^*larvae. Data points are measurements from individual larvae. n ≥ 15 larvae per group. **(E)** Quantification of ET length in NIC and *dab2* KD 6 dpf *Gt(cltca-citrine)^ct116a^*; *TgKI(dab2-mScarlet)^pd1292^* larvae. Data points are individual ETs. n ≥ 10 larvae per group. For panels C-E, error bars are SD, and P values were calculated using Welch’s t test (C), an unpaired t test (D), or a Mann-Whitney test (E). **(F-G)** Confocal images of 6 dpf *TgKI(GFP-dnm2a)^pd1274^*; *TgKI(dab2-mScarlet)^pd1292^* NIC (A) and *dab2* KD (B) LREs stained with Phalloidin-AF647 (Cyan). White arrows point to Dab2-mScarlet labeled ETs that closely associate with GFP-Dnm2a in NIC LREs (F), and white arrowheads point GFP-Dnm2a puncta assembled on enlarged ETs in *dab2* KD LREs(G). Cyan boxes are ROIs for magnified insets. **(H)** GFP-Dnm2a fluorescence intensity profile along the sub-apical region beneath the BB and perpendicular to the apical surface for 6 dpf *TgKI(GFP-dnm2a)^pd1274^*; *TgKI(dab2-mScarlet)^pd1292^* NIC (red) and *dab2* KD (blue) larvae. Each line represents a mean of maximum intensity projections of individual ROIs for an animal. n ≥ 13 larvae. **(I)** Quantification of GFP-Dnm2a puncta per cross-section from 6 dpf *TgKI(GFP-dnm2a)^pd1274^*; *TgKI(dab2-mScarlet)^pd1292^* NIC and *dab2* KD larvae. n ≥ 20 larvae. Error bars are SD. P value was calculated using a Mann-Whitney test. **(J)** Quantitation of fluorescence intensity of GFP-Dnm2a at ETs in NIC and *dab2* KD 6 dpf *TgKI(GFP-dnm2a)^pd1274^*; *TgKI(dab2-mScarlet)^pd1292^* larvae. Data points are mean values of all ROIs for individual larvae. n = 22 larvae per group. Error bars are SD. P value was calculated using a Mann-Whitney test. **(K-P)** TEM images of LREs from 6 dpf WT sibling (K, L, M) and *dab2^pd1162^* mutant larvae (N, O, P). White arrows show ETs in WT larvae (K, L, M), white arrowheads show distended ETs in *dab2^pd1162^* mutants (N, O, P). **(Q)** Quantitation of maximum ET diameter in WT and *dab2^pd1162^* mutant larvae. Data points are mean values of all ETs from individual animal. n = 8 larvae per group. Error bars are SD. P value was calculated using Welch’s t test

Because ET elongation in LREs was also caused by dynamin inhibition (Fig. 4F-H), we hypothesized that loss of Dab2 might alter Dynamin localization. To test this, we performed *dab2* KD in *TgKI(GFP-dnm2a)^pd1274^* larvae and analyzed GFP-Dnm2a distribution along ETs. Analyzing the intensity profile of GFP-Dnm2a in sub-apical region of LREs with respect to brush border using an automated image analysis pipeline, we observed that GFP-Dnm2a, which displayed a uniform distribution within the ET region in WT control larvae, instead accumulated in dense high intensity puncta on ETs (blue peaks in the graph. Fig. 6H) upon *dab2* KD (Fig. 6F-J). To investigate this phenotype further, we performed TEM analysis of 6 dpf *dab2^pd1162^*mutants and WT sibling larvae and found that loss of Dab2 caused the terminal ends of ETs to become highly distended, nearly doubling their diameter compared to those of WT LREs (Fig. 6K-Q). Together, our results suggest that Dab2 is required for the proper dynamin distribution along ETs. Furthermore, Dab2 may also regulate dynamin activity as the GFP-Dnm2a clusters seen upon loss of Dab2 in LREs may represent hyper-polymerized collars with reduced dynamin activity (35, 60). Together, our results suggest that, in addition to functioning as an endocytic adaptor, Dab2 is also a key regulator of ET morphogenesis by controlling recruitment of Myo6 and activity of Dynamin.

Next, to investigate how loss of Myo6 and Dab2 affects cargo recruitment to endocytic structures, performed mCherry gavage assays in larvae briefly pre-fixed with 4% PFA to block internalization. Unlike WT LREs, in which mCherry becomes enriched in ETs (Fig. S4C, F), *myo6a* KD larvae showed cargo accumulation at the apical membrane (Fig, S4D, F), consistent with the notion that ETs fail to form and are replaced by CCPs in the absence of Myo6. Surprisingly, although ETs are present in *dab2* mutants, mCherry cargo fails to accumulate there in *dab2* KD larvae and instead also localizes at the apical membrane (Fig. S4E, F), suggesting that Dab2 may promote recruitment of the Cubn-Amn endocytic receptor complex to ETs. Importantly, resulting from impaired ET biogenesis and altered recruitment of endocytic factors to ETs, *dab2* and *myo6* KD larvae exhibit reduced kinetics of cargo deliver to the endosomes as compared to control larvae (Fig. S4G-J).

Altogether, our results show that clathrin-coated ETs mediate fast and high-capacity protein uptake in LREs.

## Discussion

Here, we have uncovered the cellular mechanism that facilities the rapid absorption of dietary proteins by LREs. We found that LREs possess long (> 1 µm) clathrin-coated endocytic tubules (ETs). These unique structures allow LREs to internalize high volumes of protein cargoes within seconds. Protein absorption through ETs is both clathrin- and dynamin-dependent and, unlike canonical CCPs, is able to sustain the rapid kinetics of endocytosis characteristic of LREs by pre-assembling endocytic complexes along ETs. Taken together, our study suggests that continuous pinching from a pre-formed ET facilitates fast and high-capacity CME (Graphical abstract).

### LRE endocytosis as a specialized form of rapid, high-capacity CME

LREs use a form of tubular CME that shares components with canonical CCPs but is uniquely adapted to the architecture of the apical membrane of enterocytes to provide a rapid, high-capacity protein uptake. Another form of rapid CME occurs at synapses and can be completed within seconds (61), up to 20-fold faster than canonical CME (62). This fast CME activity is attributed to a priming step in which clathrin is poised near the synaptic membrane to associate with fusing vesicles during secretion (32), allowing clathrin recruitment and membrane deformation to be bypassed. Our data indicate that ETs use an analogous process that is mechanistically distinct from CME at synapses. Our TEM (Fig. 3E-L) and endogenous reporters (Fig. 3R, Fig. 4E) show that ETs are labeled with Clathrin and adaptors along their length and filled with cargo. Shedding of vesicles from the tips of ETs (Fig. 3F-H) is thus akin to an assembly line process that allows LRE endocytosis to overcome the rate-limiting steps of clathrin assembly and adaptor recruitment. Moreover, assembly of multiple structures within the same ET, suggested by TEM images (Fig. 3E-L), may also contribute to the high endocytic capacity of ETs.

In addition to allowing for fast kinetics, ETs may also facilitate cargo enrichment. Dab2, which has a central role in LRE endocytosis (13), is not only a clathrin adaptor (47) but also directly binds the Cubn-Amn endocytic receptor complex (63). This arrangement may allow ETs to be highly enriched with endocytic receptors to promote efficient cargo capture as illustrated by pHluorin uptake data showing vastly greater concentration in ETs compared to the gut lumen (Fig. 2C). In agreement, when Dab2 is lost, mCherry becomes enriched at the brush border membrane after gavage and no longer localizes to ETs (Fig. S4B), suggesting that Dab2 segregates the Cubn-Amn (CubAm) receptor complex to ETs. Together, these findings indicate that an important function of ETs is to trap and concentrate protein cargoes to facilitate their rapid uptake.

Another key effector of LRE endocytosis is Myo6, which is essential for ET formation and protein uptake (Fig. 5N-O). While loss of ETs significantly increased CCPs upon *myo6* KD (Fig. 5P), this could not compensate for the lack of ETs to maintain endocytosis in LREs (Fig. 5D-I). We also found that when ETs are replaced with CCPs, LRE microvilli length is increased (Fig. 5Q), suggesting membrane competition between the two structures. Similarly, blocking endocytosis in mouse enterocytes can result in aberrant invaginations that form at the expense of microvilli length (64), and the rate of CME also impacts microvilli length (65). These findings suggest that ETs compete with microvilli for the limited plasma membrane pool needed to support both structures. Myo6 is the only myosin family member with minus end-directed motor activity (51). Because apical F-actin is highly polarized in enterocytes, with minus ends projecting basally from microvilli tips towards the cytoplasm (53), we propose that Myo6 couples F-actin to endocytic pits and its motor activity promotes ET elongation. This model is supported by our data showing F-actin bundles projecting beneath the apical membrane alongside ETs and our TEM analysis showing long microfilaments residing near ETs. Future studies manipulating Myo6 motor activity may allow this idea to be tested directly. Alternatively, Myo6 may spatially restrict dynamin activity to the tips of ETs, preventing fission from occurring near the brush border membrane and promoting ET elongation. This could explain why endocytic structures resemble CCPs in *myo6a* KD LREs (Fig. 5N).

Dab2 also has a role in ET biogenesis. This could in part be mediated by Dab2-dependent recruitment of Myo6 to ETs (Fig. 5B, C). However, unlike *myo6* KD, loss of Dab2 results in elongated and highly distended ETs. This may be due to an effect of Dab2 on Dynamin activity at ETs, which is supported by our data showing that loss of Dab2 increases ET length similarly to the effect of dynamin inhibition (Fig. 6B, E, Fig. 4F-H), and by the formation of Dnm2 clusters on ETs (Fig. 6G). These structures are similar in appearance to hyper-polymerized dynamin collars that form in the absence of GTP to inhibit vesicle fission (35, 66). However, it is unclear how Dab2 could regulate the GTPase activity of Dnm2. Unlike most GTPases that require other proteins to control nucleotide hydrolysis and exchange, dynamins are autoregulatory. However, BAR domain proteins, which directly bind dynamin, have been proposed to influence its GTPase activity (67). Because Dab2 binds to some BAR domain proteins like Fcho2 (68), it is possible that additional effectors link Dab2 to the regulation of dynamin activity. Future studies investigating if these interactions are important for regulating dynamin activity in LREs may help elucidate this intriguing phenotype.

### ETs as a conserved mechanism for high-capacity endocytosis

Long apical invaginations have been observed in TEM studies of rat proximal tubule cells (69), and they were found to harbor clathrin coats (24). While these endocytic structures have not been studied in detail, they have been proposed to facilitate protein reabsorption in the kidney during high demand, when CCPs can become saturated (70). Because Dab2 and clathrin both showed high endogenous expression in the pronephros (Fig. 3A, B), the larval kidney in zebrafish, we investigated if these cells possess ETs. We found strong morphological and molecular similarity to LREs, with overlap of Myo6 with Dab2-mScarlet on tubular invaginations labeled by Cltca-Citrine and adjacent to F-actin bundles (Fig. S3B). These findings suggest that ETs are not unique to LREs but may be a common feature of epithelial cells specialized for apical endocytosis.

ETs also bear striking resemblance to endocytic structures found in yeast. During yeast endocytosis, actin polymerization and myosin activity drive elongation of endocytic pits into deep invaginations (71–73) that are capped with clathrin (29). Actin polymerization (29) and myosin activity (73, 74) are required for pit invagination and endocytosis in yeast. By contrast, actin and myosin are not thought to be directly involved in endocytosis in mammalian cells in culture (75), except in cells with increased membrane tension (76, 77). Our in vivo studies in zebrafish here suggest that myosin-dependent membrane invagination originally discovered in yeast is in fact a highly conserved process needed for high-capacity endocytosis.

Another intriguing similarity between endocytosis in yeast and LREs is that both require phosphoinositide phosphatase activity. In yeast, turnover of PI(4,5)P2 by the phosphatase Sjl2p is required for endocytosis (78) and is thought to spatially restrict PI(4,5)P2 to promote membrane curvature that is needed for vesicle scission (79). Although vesicle scission is dynamin-dependent in LREs, the PI(4,5)P2 phosphatase Tpte (Vsp) was shown to be required for endocytosis in LREs (80, 81), suggesting it may have similar roles in membrane curvature. Future studies investigating how PI(4,5)P2 segregation relates to ET biogenesis will further shed light on mechanistic similarities of endocytosis in yeast and LREs.

Altogether, this work studying the function of LREs in zebrafish uncovered a unique form of CME that occurs through long preformed tubules (ETs). Our data support a model in which ETs can overcome the rate-limiting steps of membrane deformation and nucleation during CME, allowing them to continuously generate endocytic vesicles from cargo-filled ETs (Graphical abstract). The remarkable cargo handling capacity of ETs is underscored by in vivo kinetic assays that illustrate how a relatively small population of LREs facilitates dietary protein absorption and organismal growth. Our study highlights the importance of investigating endocytosis in professional absorptive cells in vivo and suggests that ETs may represent a conserved mechanism for high-capacity uptake.

## Author contributions

Conceptualization: I.M., M.B., D.S.L.; Methodology: I.M, D.S.L., M.B., T.R.L., C.C.; Validation: I.M., D.S.L., C.C.; Formal analysis: I.M., D.S.L., M.B., C.L.B., K.L.; Investigation: I.M., D.S.L., T.R.L., C.C., C.L.B., S.W., J.P.; Software: K.L.; Resources: MS.B., T.R.L., V.Y.A.; Writing - original draft: I.M., D.S.L., M.B.; Writing - review & editing: M.B., D.S.L., V.Y.A., T.R.L.; Visualization: D.S.L.; Supervision: M.B., D.S.L., V.Y.A.; Project administration: M.B., D.S.L.; Funding acquisition: M.B., V.Y.A.

## Acknowledgements

We thank Ricardo Vancini, Stella Finkelstein and Ying Hao for their help with electron microscopy, The Duke Zebrafish Core for fish care, and the Duke Light microscopy core facility for equipment access. We also thank Kalina Hristova for fluorescent protein plasmids and Le Trinh for providing the *Gt(cltca-citrine)^ct116a^* line. We would like to thank Terry Lechler and John Rawls for critical reading of the manuscript and the members of the Bagnat and Rawls lab for discussions. This work was funded by National Institute of Diabetes and Digestive and Kidney Diseases (NIDDK) grant R01DK137812 to MB, and by National Eye Institute P30 EY005722 to VYA. TRL was supported by National Eye Institute grant EY033763; CB was supported by NIDDK grant T32DK007568.

## Competing interests

The authors declare no competing interests.

## Materials and methods

### Fish

Zebrafish were maintained and used in accordance with the guidelines of the Duke University Institutional Animal Care and Use Committee (IACUC). Zebrafish (*Danio rerio*) stocks were maintained at 28°C in a recirculating aquatic system. Embryos were obtained by natural spawning, with one male and one female placed together in a mating tank under controlled light–dark cycle conditions (82). Genotypes were determined by fin clipping. Male and female breeders between 3 - 18 months of age were used to generate experimental animals. Zebrafish larvae aged 1-6 days post-fertilization (dpf) from the Ekkwill (EK) or AB/TL genetic backgrounds were used for all experiments. Sex-based effects were not evaluated because sex determination in zebrafish occurs during the late juvenile stage, well beyond the developmental window examined in this study. Strains generated for this study are: *clta^pd1247^, TgKI(dab2-mScarlet)^pd1292^, TgKI(GFP-dnm2a)^pd1274^* and *TgKI(Slc10a2-mScarlet)^pd1308^*. Strains used in this study and published previously are: *dab2^pd1162^* (13), *Tg(hsp70l:p75-GFP)^pd1180^* (39)*, Tg(hsp:Venus-FYVE)^pd1046^* (83), *Gt(cltca-citrine)^ct116a^* (46) and *TgKI(GFP-rab11a)^pd124^* (84).

### Mice

All experiments were performed with Institutional Animal Care and Use Committee approval from Duke University, and in accordance with the guidelines of the Division of Laboratory Animal Resources guidelines for the humane treatment of animals. Mouse lines used in this study were: wild type (C57BL/6J; JAX:000664, RRID:IMSR_JAX:000664). Both male and female mice were used. Mice were maintained in a facility with a 12 h light/dark cycle with *ad libitum* access to food and water.

### Methods Details

#### Genome editing

Stable Mutants, endogenous reporter knock-in lines, and transient knockdown were generated using CRISPR/Cas9. CRISPRscan (85) was used to identify gRNA target sites, while oligo-based method (86) was used for gRNA design and synthesis using HiScribe T7 Transcription Kit (NEB E2050S). For generating the c*lta^pd1247^*mutants, one-cell stage EK embryos were injected with 500 ng/μl Cas9 protein (PNA bio CP02) and 50 ng/μL of gRNA (gRNA target sequence: 5’ GAAGCGCAGGGAACGGTGTCGGG 3’). For the c*lta^pd1247^* mutants, gRNA was designed to target exon 1, immediately downstream of the start codon (Fig. S2 A). The mutant allele has a 10 bp insertion at +51, leading to a frame shift after amino acid 17 and a premature stop codon encountered after amino acid 44 (Fig. S2B, C). Genotyping for c*lta^pd1247^* mutant was performed using 5’-GCACATTCAGCAAGCCATGGA-3’ forward and 5’-GAGAATGACGAGGGCTTCAG-3’ reverse primer.

For transient knockdown (KD) of *cltb, myo6a* and *dab2,* multiple regions of the gene were targeted using CRISPR/Cas9*. For cltb* KD, two gRNA targeting exon 1 (5’-TACTAACGTTTGAGCTGT-3’ and 5’-TTTTTCCGCCTTCGACAA-3’), one targeting exon 3 (5’-AAAGTCTGCGCAAGTGGA-3’) and one targeting exon 6 (5’-ACTCTGCGATTTCAACCC-3’) were used. For *dab2* KD, 4 guide RNAs targeting exon 1 (5’-GCATCGACCACTGAATCA-3’), exon 2 (5’-TCCTCTTGGCCAGATTTC-3’), exon 3 (5’-AATAGCTGCTCGTTCTCA-3’) and exon 4 (5’-CACTGATAACCGGGCATT-3’) where used*. myo6a* was targeted using 4 gRNA against intron proceeding exon 1 (5’-GGCGTGTGTCGGAGGATG-3’), exon 1 (5’-CACCCTACAGACGGGTT-3’), exon 12 (5’-CTTCGACTCGTCTGCGCA-3’) and exon 34 (5’-GGGGGAATCCAGTACCTG-3’). Pooled synthesis was performed using HiScribe T7 Transcription Kit (NEB E2050S) as described above except for pooling the annealed gRNA templates before the T7 transcription reaction. 300 ng/μL, 50 ng/μL, 150 ng/μL of *cltb*, *dab2* and *myo6a* gRNA pool was co-injected with 500 ng/μL Cas9 protein to obtain knockdown of the respective genes.

#### Transgenic Knock-in zebrafish

For generating endogenously tagged knock-in lines, we used our strategy we previously described (84). The composition of the injection mixes for all targets were: PCR donor (20 nM), gRNA (4 µM), Cas9 protein (3 µM) (PNA Bio CP-02), and Phenol Red (0.05%).

##### TgKI(dab2-mScarlet)^pd1292^

A gene fragment spanning from the middle of intron 14 to the last coding sequence codon of exon 15 of *dab2* was cloned into pUC19-TgKI-MCS-tdTomato-polyA (Addgene Plasmid # 174022) using cut-and-paste cloning with the following primers: dab2-I14-F: 5’-ATGCCCGCGGGAGTTCGGACGTTTCATAGAGTTG-3’ and dab2-E15-R: 5’-ATGCCTCGAGGGCAAAGGGATTTCCCCCAAA-3’.

The TdTomato sequence was then replaced with mScarlet using In-Fusion seamless cloning (Takara Bio) with the following primers: mScarlet-F: 5’-CCCTTTGCCCTCGAGGTGAGCAAGGGCGAGGCAG-3’ and mScarlet-R: 5’-ACTGAGAATGATTTACTTGTACAGCTCGTCCATGCCG-3’. The gRNA target site used for *dab2* was: dab2_gRNA_site: TGTACTGTGTAGCCATGGGCCGG.

##### TgKI(Slc10a2-mScarlet)^pd1308^

A gene fragment spanning from the middle of intron 4 to the last coding sequence codon of exon 5 of *slc10a2* was cloned into pUC19-TgKI-MCS-mScarlet-MCS-polyA (Addgene Plasmid #174023) using In-Fusion seamless cloning (Takara Bio) with the following primers: slc10a2-I4-F: 5’-TTCGCTAGCCCGCGGCTATGACCGGGATACAGTCAG-3’ and slc10a2-E5-R: 5’-CACCTCGAGTCTAGACAATTTTGTACCTTTGTCTTTGTTTTG-3’.

The gRNA sites used for genomic targeting were then mutated in the donor plasmid using Q5 site-directed mutagenesis (New England Biolabs, NEB) with the following primers: slc10a2-SDM-F: 5’-CCCAAATGAACCACAGATGTGAAAGTAGC-3’ and slc10a2-SDM-R: 5’-CCAGCGTGAGATCAACAAATGATACACTGTAG-3’. The gRNA target site used for *slc10a2* was: slc10a2_gRNA_site: GTTCATTTGGGCCAGAGCAAGGG.

##### TgKI(GFP-dnm2a)^pd1274^

A gene fragment spanning from the 5’ UTR to the middle of intron 1 of *dnm2a* with GFP placed before the start codon was cloned into pCS2+ with a multi-site In-Fusion reaction (Takara Bio) using the following primers: dnm2a-5’-F: 5’-TCGAATTCAAGGCCTCTCGACATGTGCTGGTTAAGTTGGTG-3’, dnm2a-5’-R: 5’-TGCTCACCATGGTTTCTGGCCGTCTCTCAG-3’,GFP-F: 5’-GCCAGAAACCATGGTGAGCAAGGGCGAGG-3’, GFP-R: 5’-GGTTGCCCATCTTGTACAGCTCGTCCATGCC-3’, dnm2a-3’-F: 5’-GCTGTACAAGATGGGCAACCGGGGGATG-3’, dnm2a-3’-R1: 5’-ACGACTCACTATAGTTCTAGCATGACGTCAATGTCCAGAGAG-3’.

The gRNA sites used for genomic targeting were then mutated in the donor plasmid using Q5 site-directed mutagenesis (New England Biolabs, NEB) with the following primers: dnm2a-SDM-F: 5’-GAGCCCGTTCCTGCGCGCTGAGAGACGGC-3’ and dnm2a-SDM-R: 5’-GCAGCGTCCATACGCTCGGGTTACTTTCAAGTGTTCAG-3’. The gRNA target site used for *dnm2a* was: dnm2a_gRNA_site: GGAACGGGCTCGCAGCAGCAGGG.

#### Soluble fluorescent protein production

pRSET-B-mCherry and pRSET-B–GFP2 vectors were gifts from Kalina Hristova (87) while pRSET-pHluorin-tagRFPt vector was made during this study. The mCherry sequence in pRSET-mCherry present between the BamH1-EcoR1 restriction enzyme sites was swapped with PCR amplified pHluorin-tagRFPt fragment generated from pME:B4GALT1-pHluorin-tagRFPt (39) using 5’-ACTGGATCCTCCGGTCATGAGTAAAGGAGAAGA-3’ forward and 5’-ACTGAATTCTTACTTGTACAGCTCGTCCATGCC-3’ reverse primers containing BamH1 and EcoR1 overhangs respectively. mCherry, GFP and pHluorin-tagRFPt soluble fluorescent proteins were produced by uninduced expression in the BL21-Gold (DE3) strain of Escherichia coli (87). Cells were lysed in Bacterial lysis buffer (300 mM NaCl, 10 mM MgCl2, 0.1% NP-40, 50 mM Tris-Cl (pH 8.0), 0.5% Glycerol) supplemented with Lysozyme (final = 200 µg/mL), PIC and PMSF (1mM) followed by 3 rounds of sonication. The lysates were centrifuged at 21130 g for 25 min after allowing for 30 minutes of lysis on ice. The fluorescent proteins were purified from the soluble bacterial cell lysates using His tag affinity purification using Ni-NTA beads (Invitrogen, Cat. #R90115). The purified fluorescent proteins were eluted in 1X PBS containing 250 mM imidazole followed by buffer exchange with 1X PBS to reduce final imidazole concentration below 750 μM, which is below the tolerable limit for mammalian cells (2 mM).

#### Microgavage

Gavage experiments of 6 dpf larvae were performed as previously described (88). Larvae were gavaged with ∼65 nL of 0.5 mg/mL mCherry, 2 mg/mL GFP, 2 mg/mL pHluorin-tagRFPt, 200 μM FM4-64 (Thermo Scientific, F34653), 0.9 mg/mL Sulfo-NHS-SS-Biotin, 0.4 mg/mL Streptavidin-AF647, 2 mg/mL Lucifer Yellow CH, Lithium Salt (Invitrogen, L453) or 50 μg/mL Wheat Germ Agglutinin WGA, Alexa Fluor 594 (ThermoFisher, W11262). mCherry, GFP, Phluorin-tagRFPt stocks were made in 1X PBS, FM4-64 and WGA stocks were made in ultrapure H2O and all working gavage solutions were diluted in 1X PBS.

#### Cell surface biotinylation

To label the luminal membrane of the brush border, larvae were gavaged with 0.9 mg/mL of Sulfo-NHS-SS-Biotin dissolved in PBS, incubated for 15 minutes on ice, and then gavaged with excess 1X PBS in 10 mM Tris, pH 8. Next, larvae were gavaged with 0.4 mg/mL of Streptavidin conjugated to Alexa Fluor 647 dissolved in 1X PBS and incubated for 5 minutes on ice. Finally, larvae were gavaged with excess 1X PBS and incubated at room temperature for various time points before fixation according to the duration of the chase periods that are described in the figures and legends.

#### Cargo uptake and kinetics assay

For assaying total uptake in LREs, 0.5 mg/mL mCherry (Fig. 4I-S, Fig. S4E) or a mix of 0.5 mg/mL mCherry and 2 mg/mL Lucifer yellow (Fig. 5D-G) were gavaged in 6 dpf WT zebrafish larvae and fixed with 4% PFA 4 hours post gavage.

Early kinetics of cargo endocytosis and plasma membrane turnover in 6 dpf zebrafish AECs and LREs was probed by fast fixation of gut cells by 4% PFA gavage before fixing the whole larvae in 4% PFA at the desired early time points following 2 mg/mL GFP +200 μM FM4-64 gavage (Fig. 1B-F) in WT, 2 mg/mL GFP in *TgKI(Slc10a2-mScarlet)^pd1308^* (Fig. 1I-L) or 0.5 mg/mL mCherry in *Tg(hsp70l:p75-GFP)^pd1180^* (Fig. S1 A-F).

For completely halting endocytic activity, 6 dpf zebrafish larvae were pre-fixing by 4% PFA gavage for 30 seconds before GFP + WGA-AF594 (Fig. 1 N-P) or Sulfo-NHS-SS-Biotin (Fig. 3Y, Z) or mCherry (Fig. 3V, W) test cargo gavage. Following test cargo gavage, the whole larvae were fixed at desired time points in 4% PFA overnight.

#### Dyngo4a inhibitor treatment

For inhibiting Dynamin activity Dyngo4a (Hydroxy Dynasore) (Tocris, C5364) 1 mM stock solution was made in DMSO while working gavage solution, 40 μM and 100 μM Dyngo4a was made by diluting in 1X PBS. Zebrafish larvae gavaged with equivalent concentration of DMSO diluted in 1X PBS were used as controls.

#### Immunohistochemistry and Fluorescence microscopy of zebrafish samples

For high resolution imaging, 4% PFA fixed zebrafish larvae were washed 2 x 5 minutes with 1X PBS and embedded in 5% low-melting Agarose, cured for 30 minutes at 4° C and then sectioned using a vibratome (VT 1000S; Leica). Next, 170 μm sections were collected and stained with DAPI (ThermoFisher, 62248) (∼200 nM) and Phalloidin-AF647 (ThermoFisher, A30107) (∼33 nM) (Fig. 1 C-F, I-L, O, P) or Phalloidin-AF594 (∼33 nM) (ThermoFisher, A12381) alone (Fig. 3Q-R, Z) or Phalloidin-AF647 (∼33 nM) (Fig. 3V-W, Fig. 4D, Fig. 6F, G, Fig. S1A-F, Fig. S2D-E, Fig. S4A-C) diluted in 1X PBS followed by 5 × 10 minutes washes with 1X PBS. The sections were then mounted on a glass slide using mounting medium containing 20 mM Tris (pH 8.0), 90% glycerol, and 0.5% *n*-propyl gallate. For phalloidin-AF647 and Rabbit α-Myosin6 (primary antibody, Proteus, Cat. # 25-6791, 1:300) and goat α-Rabbit AF405 (secondary antibody, ThermoFisher, Cat. # A48254) (Fig. 5B-C, J, K, Fig. S3 B) or phalloidin-AF405 (∼33 nM) and Rabbit α-Myosin6 (primary antibody, Proteus, Cat. # 25-6791, 1:300) and goat α-Rabbit AF647 (secondary antibody, ThermoFisher, Cat. # A-21245, 1:500) (Fig. 6A-B) or phalloidin-AF647 and Donkey α-Rabbit AF488 (secondary antibody, ThermoFisher, Cat. #A-21206), the sections were first incubated with blocking solution [5% Bovine Serum Albumin (Sigma Aldrich, Cat. # A7906) in PBS containing 0.5% Triton-X100 (0.5% PBSTx) (Sigma Aldrich, Cat.# X100)] for 1-5 hours followed by primary antibody incubation in blocking solution for 1 hour at room temperature or overnight at 4° C followed by 5 × 10 minutes washes with 0.1% PBSTx and secondary antibody incubation in blocking solution and subsequent washes with 0.1% PBSTx. Sections were finally washed 2 x 5 minutes with 1X PBS before mounting on glass slides as described above. Images were collected using either a Leica SP8 microscope equipped with 63X/1.4NA objective or Olympus (Evident Scientific) FV4000 microscope equipped with 60X/1.5NA objective.

Overlap of Cltca-citrine and gavaged mCherry was done in 6 dpf zebrafish *Gt(cltca-citrine)^ct116a^*. Larvae were gavage with 0.5 mg/mL mCherry followed by flushing by 1X PBS gavage after 15 minutes of mCherry gavage. Live fish were mounted in 1.3% low-melting agarose and imaged on Leica SP8 microscope using 10X objective and Galvano scanner in tile scan mode. Adjacent tiles were stitched post-acquisition using Leica Application Suite X (LAS X) software environment, utilizing the integrated Mosaic / Tile Scan stitching module to generate the final image (Fig. 3P). Similarly, whole larvae live images were generated for *TgKI(dab2-mScarlet)^pd1292^*; *Gt(cltca-citrine)^ct116a^* (Fig. 4 A, B) and *TgKI(GFP-dnm2a)^pd1274^*(Fig. 4 C).

For assaying total uptake in LREs (Fig. 4J-L, P-S, Fig. 5D-G, Fig. S4E), 6 dpf larvae were fixed after gavage and were washed with 2 x for 5 minutes with 1X PBS, mounted in 1.3% low-melting agarose on a glass-bottom imaging dish, and imaged either on a Leica SP8 microscope using a 25X objective or on a Olympus FV4000 microscope using a 30X objective.

#### Immunohistochemistry and Fluorescent microscopy of mice samples

Mice were euthanized by rapid decapitation at postnatal day 6 (P6) or carbon dioxide inhalation at P36. Intestines were collected, and the middle region of the ileum was dissected. Samples were washed in 1X PBS and fixed overnight in 4% PFA at 4° C. Intestinal segments were subsequently embedded in 5% low-melting-point agarose prepared in PBS and sectioned at 125 μm using a vibratome (Leica VT1000).

For immunofluorescence staining, sections were washed three times in 1X PBS and blocked for 1 hour in 5% normal goat serum (NGS) and 3% bovine serum albumin (BSA) prepared in PBSTx. Sections were then incubated overnight at 4° C with an antibody against clathrin light chain A (CLTA; 1:200; BioLegend, 811501) diluted in blocking buffer. The following day, sections were washed three times in PBSTx and incubated with Alexa Fluor 488-conjugated goat anti-mouse secondary antibody (1:500; Invitrogen, A11001) diluted in blocking buffer. After 90 minutes, phalloidin (1:100; Invitrogen, A22287) was added for an additional 30 minutes, resulting in a total secondary antibody incubation of 2 hours. DAPI (1:100,000; Invitrogen, D1306) was added during the final 10 minutes of incubation. Sections were then washed three times in 1X PBS, mounted using mounting medium containing 20 mM Tris (pH 8.0), 90% glycerol, and 0.5% *n*-propyl gallate. High-resolution images were acquired using an Olympus FV3000 confocal laser-scanning microscope equipped with a 60X objective and 3X digital zoom. Images were acquired at a resolution of 2048 × 2048 pixels with a z-step size of 0.32 μm. Five to eight images were acquired per animal from multiple fields of view and intestinal sections. Images were subsequently deconvolved using Huygens Professional software (version 19.10.0p3, 64-bit).

#### Live imaging

Live imaging of pHluorin-TagRFPt gavaged 6 dpf larvae was performed within 10-20 minutes of gavage by mounting the live larvae in 0.9% low-melting agarose in a glass bottom imaging dish and imaged using 25X Leica objective on the Leica SP8 confocal microscope (Fig. 2 C, N).

#### *In Situ* Hybridization Chain Reaction (HCR)

HCR probes for *myo6a*, hairpins, amplification, wash, and hybridization buffers were purchased from Molecular Instruments (56). HCR was performed as described previously (37).

#### Transmission Electron Microscopy

6 dpf zebrafish larvae were fixed in 2.5% Glutaraldehyde in 0.1 M Na-cacodylate at room temperature for 3 hours. Samples were then washed in 0.1 M sodium cacodylate buffer (3 changes for 15 minutes each). Samples were then post-fixed in 1.0% OsO4 in 0.1 M Sodium cacodylate buffer for 1 hour on rotator. After that samples were washed in 3 changes of 0.1M sodium cacodylate, washing for 15 minutes between each change. Samples were then placed into *en bloc* stain (1% uranyl acetate) overnight at 4° C. Subsequently they were dehydrated in a series of ascending acetone concentrations including 50%, 70%, 95%, and 100% for three cycles with 15 minutes incubation at each concentration change. Samples were then placed in a 50:50 mixture of Epon:acetone overnight on a rotator. This solution was then replaced twice with 100% epoxy resin (epon) for at least 2 hours at room temperature on a rotator. Samples were embedded with 100% epon resin in beem capsules for 48 hours at 60° C. Samples were then ultrathin sectioned to 60-70 nm on a Reichert Ultracut E ultramicrotome. Harvested grids were then stained with 2% uranyl acetate in 50% ethanol for 30 minutes and Sato’s lead stain for 1 minute. Images were acquired using a Phillips CM12 electron microscope operating at 80Kv (Fig. 3A-L, Fig. 6K-P).

To analyze ET morphology upon *myo6a* KD, 6 dpf zebrafish larvae were fixed in 2.5% glutaraldehyde (in 0.1 M Na-cacodylate) at room temperature for 3 hours. Samples were then washed in 0.1 M sodium cacodylate buffer (3 changes for 5 minutes each). Samples were then post-fixed in 1% tannic acid in 0.1M HEPES (pH7.4) for 1 hour on rotator. After that samples were washed in 3 changes of 0.1 M maleate buffer, pH 6, washing for 5 minutes between each change. Samples were then placed into *en bloc* stain (1% uranyl acetate in maleate buffer) for 1 hour followed by rinse with maleate buffer, pH 6, 2 x 3 minutes. Subsequently they were dehydrated in a series of ascending ethanol concentrations including 50%, 70%, 80%, 90% and 95% for three cycles with 15 minutes incubation at each concentration change followed by 3 changes of 100% ethanol at an interval of 30 minutes. Samples were then infiltrated with Spurr resin with a series of 3:1,1:1 and 1:3 mix of Ethanol/Spurr with interval of 30 minutes between changes before transferring to 100% Spurr where they were kept overnight. Samples were embedded by curing at 65°C for 2 days. Samples were then ultrathin sectioned to 60-70 nm on a Leica EM UC6 ultramicrotome and collected on copper grids. Grids were then stained with 2% uranyl acetate for 30 minutes and 3.5% lead citrate for 1 min. Images were acquired with a JEM-1400 electron microscope (JEOL) at 60 kV with a digital camera (BioSprint; AMT) (Fig. 5M, N).

#### RT-PCR

Total RNA was extracted from 6 dpf zebrafish larvae using TRIzol reagent (Invitrogen, Cat. # 15596026) according to the manufacturer’s instructions. First-strand cDNA was synthesized from total RNA using SuperScript III Reverse Transcriptase (Invitrogen, Cat. # 18080-093) with oligo(dT) primers. PCR amplification was performed using OneTaq 2X Master Mix (NEB, Cat.# M0482L) with gene-specific RT-PCR primers, 5’-CGATGCGAACTCGCGTAAACAGGA-3’ forward and 5’-TTGTAGAAATCCTCATCCAGCACC-3’ reverse for *clta*, 5’-GAAAGTCTGCGCAAGTGGAG-3’ forward and 5’-CTCCAGCTCCTTCTTGGC-3’ reverse for *cltb* and 5’-ATCACACCTTCTACAACGAGC-3’ forward and 5’-CATCACCAGAGTCCATCACG-3’ reverse for *actb* (β-actin) which was used as a housekeeping control (89). PCR products were run on 2% agarose gel, band intensities were measured using ImageJ software and plotted using GraphPad Prism (GraphPad Software) (Fig. S2D, E).

##### Image analysis

Unless noted otherwise, all image analysis was performed using Fiji (ImageJ2, version 1.54f).

##### TEM analysis

Depth of endocytic invagination (Fig. 3M), length of microvilli (Fig. 3O, Fig. 5Q) and maximum diameter of ETs (Fig. 6Q) were measure by using the line tool, whereas the number of ETs and CCP per micron was visually counted along the length of apical surface for each image (Fig. 3N, Fig.5O, P). Darker stained microvilli clefts showing an invaginated appearance within 100 nm from the brush border were counted as CCPs whereas longer invaginations were counted as ETs.

##### Fluorescence microscopy analysis

Relative cargo uptake along the length of LREs was quantified from max projected confocal z-stack images of LRE region as described previously (13) (Fig. 4M, T, Fig. 5H, I, Fig. S4 F). Mean and standard error mean (SEM) along the length of LRE calculated.

For kinetics of GFP and FM4-64 (Fig. 1G, H) and Slc10a2-mScarlet over time (Fig. 1M) and, mCherry and p75-GFP (Fig. S1G, H), a ratio of intracellular to brush border intensity was collected by measuring the mean pixel intensity of the brush border (identified using the phalloidin channel), and an intracellular ROI of a similar area immediately below the brush border using the freehand selection tool for individual cells.

For colocalization analysis, the sub-apical region was first segmented using wand tool with high thresholding to include a wider region to obtain ROIs. Each ROI consisted of 2-5 neighboring cells. Multiple ROIs were marked from a single image. Pearson’s R value (above threshold) for co-localization was obtained for individual ROIs across multiple images by using the Coloc2 tool (Fig. 1Q, Fig. 2M, Fig. 3X).

For pH sensor gavage experiments, ET regions and endosomes within LREs were first segmented, and pHluorin and tagRFPt intensity values were measured. ETs were identified using the pHluorin channel to measure both pHluorin and tagRFPt intensities, while endosomes were identified using the tagRFPt channel to measure both pHluorin and tagRFPt intensities. Endosomes were also analyzed from time-lapse imaging of pHluorin-TagRFPt gavaged zebrafish larvae (Fig. 2D, O). A Standard curve of pH-dependent pHluorin and tagRFPt fluorescence intensity was generated using experimental data we previously published, in which we incubated pHluorin-tagRFPt in pH-calibrated phosphate-citrate buffers and then collected intensity values using pre-defined confocal imaging parameters (39) (Fig. 2B).

For obtaining Myo6 and GFP-Dnm2a intensity on ETs, ROIs were marked by segmenting ETs using Cltca-citrine signal, followed by measurement of Myo6 antibody staining (Fig. 6C) and GFP-Dnm2a (Fig. 6J) fluorescence intensity with the ROIs.

Numbers of Cltca-Citrine labeled ETs per micron were visually counted along the length of apical surface for each confocal LRE cross-section image to calculate the frequency of ETs in *dab2* kd (Fig. 6D). The number of GFP-Dnm2a puncta obtained after applying an identical threshold for all images and were counted for each LRE cross-section image (Fig. 6I). ET length upon Dynamin inhibition (Fig. 4H) and *dab2* kd (Fig. 6E) was obtained by using the line tool to trace Cltca-Citrine labeled ETs and measure individual ETs from the confocal images of LRE cross sections. Similarly, length of Dab2-mScarlet upon *myo6a* knockdown (Fig. 5L) and length of Clta in mice ileum (Fig. 3U) were measured.

As a measure of relative localization of gavaged mCherry with respect to the brush border rectangular ROIs were drawn perpendicular to the apical surface (Fig. S4D). The ROI size was kept constant and same ROI was applied to multiple cells within a cross-section image and across multiple such images to obtain mCherry and Phalloidin intensity profiles. Mean peaks distance for mCherry intensity profile was calculated by Σ(x*y)/Σy and then subtracted from the BB tip distance (specified by the first distance value after 2000 AU, which corresponds to the sharp increase in Phalloidin intensity at the tip of the BB) to find the distance between mean peaks of mCherry from the BB tip thus reflecting the relative position of mCherry with respect to BB (Fig. S4D). For obtaining the percentage of internalized mCherry in the endosomes of LREs, Venus-FYVE labeled endosomes were segmented and mCherry fluorescent intensity in these endosomes was measured for each expressing LRE. Total mCherry intensity in the corresponding LRE cell was used to calculate the percentage of mCherry in endosome for each cell (Fig. S4J).

To obtain the nature of GFP-Dnm2a intensity distribution in the ET region upon *dab2* KD, confocal LRE cross-section images of non-injected controls and *dab2* KD (Fig. 6F) were used for automated image analysis using a custom Python script [Python (Version 3.11), scikit-image (Version 0.26.0), SciPy (Version 1.17.1), and OpenCV (Version 4.13)]. Using the phalloidin stained BB as a reference, outwards facing line segments of 5 µm were drawn perpendicular to the BB at intervals of 1 µm. Equal sized rectangular ROIs of width 1 um were then centered around these line segments. GFP-Dnm2a signals within these ROIs were max projected across the long axis to generate intensity profiles and plotted for analysis (Fig. 6H).

##### Quantification and Statistical analysis

GraphPad Prism Version 11.1.0 for Mac and Microsoft Excel were used to analyze and plot data. Statistical analysis for the comparisons of invagination depth (Fig. 3M), ET length (Fig.4H, Fig. 6E), extent of co-localization of Strep-Af647 and GFP-Rab11a compared to extent of colocalization of mCherry and Venus-FYVE (Fig. 2M), ET frequency (Fig. 3N, Fig. 6D), MV length (Fig. 3O, Fig. 5Q) Dab2-mScarlet and Cltca length (Fig. 3U, Fig. 5L), pHluorin/TagRFPt ratio (Fig. 2D), relative transcript levels (Fig. S2D, E), maximum ET diameter (Fig. 6Q), number of ETs and CCPs per micron (Fig 5O, P), Myo6 and GFP-Dnm2a fluorescence intensity on ETs (Fig. 6C, J), *clta* and *cltb* mRNA expression (Fig. 3D, E), number of GFP-Dnm2a puncta (Fig. 6I) and mCherry distance from Brush border (Fig. S4D) was performed by two-tailed unpaired t-tests in GraphPad Prism.

Fluorescent cargo uptake in LREs was analyzed using a previously described method (13) where anterior to posterior intensity profile for fluorescent cargo uptake in LREs were first processed in Microsoft excel to find mean of 40 μm bins along the AP axis for individual larvae which was then used for comparing groups by Two-way ANOVA in GraphPad Software (Fig.4M, T, Fig.5 H, I, Fig. S4H).

Statistical significance for colocalization of markers (Pearson’s R value) was performed using One sample t and Wilcoxon test (Fig. 1Q, Fig. 3X).

For kinetic analysis of gavaged and genetically encoded fluorescent proteins, data were analyzed by Two-way ANOVA. The P values shown in figure panels are multiple comparisons tests for LREs or AECs at t = 0.5 minutes versus t = 3 minutes (Fig. 1G, H, M, Fig. S1G, H). The P value for Fig. S4J is a multiple comparisons test for NIC versus both *dab2* KD and *myo6* KD at t = 3 minutes.

Sample sizes for each experiment are indicated in the respective figure legends. A P value below 0.05 was considered as a significant difference between the groups.

## STAR★Methods

### Key resources table

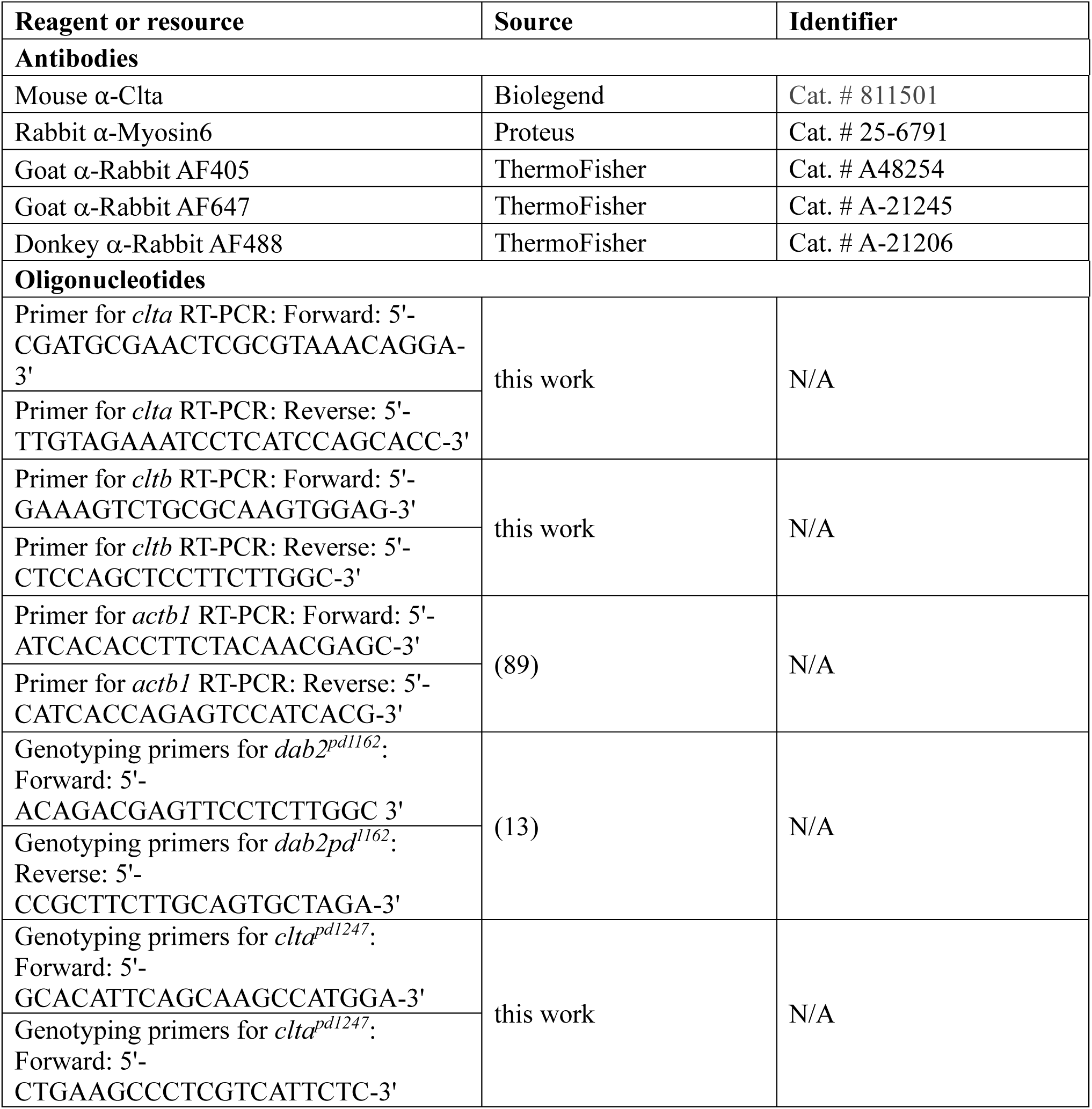
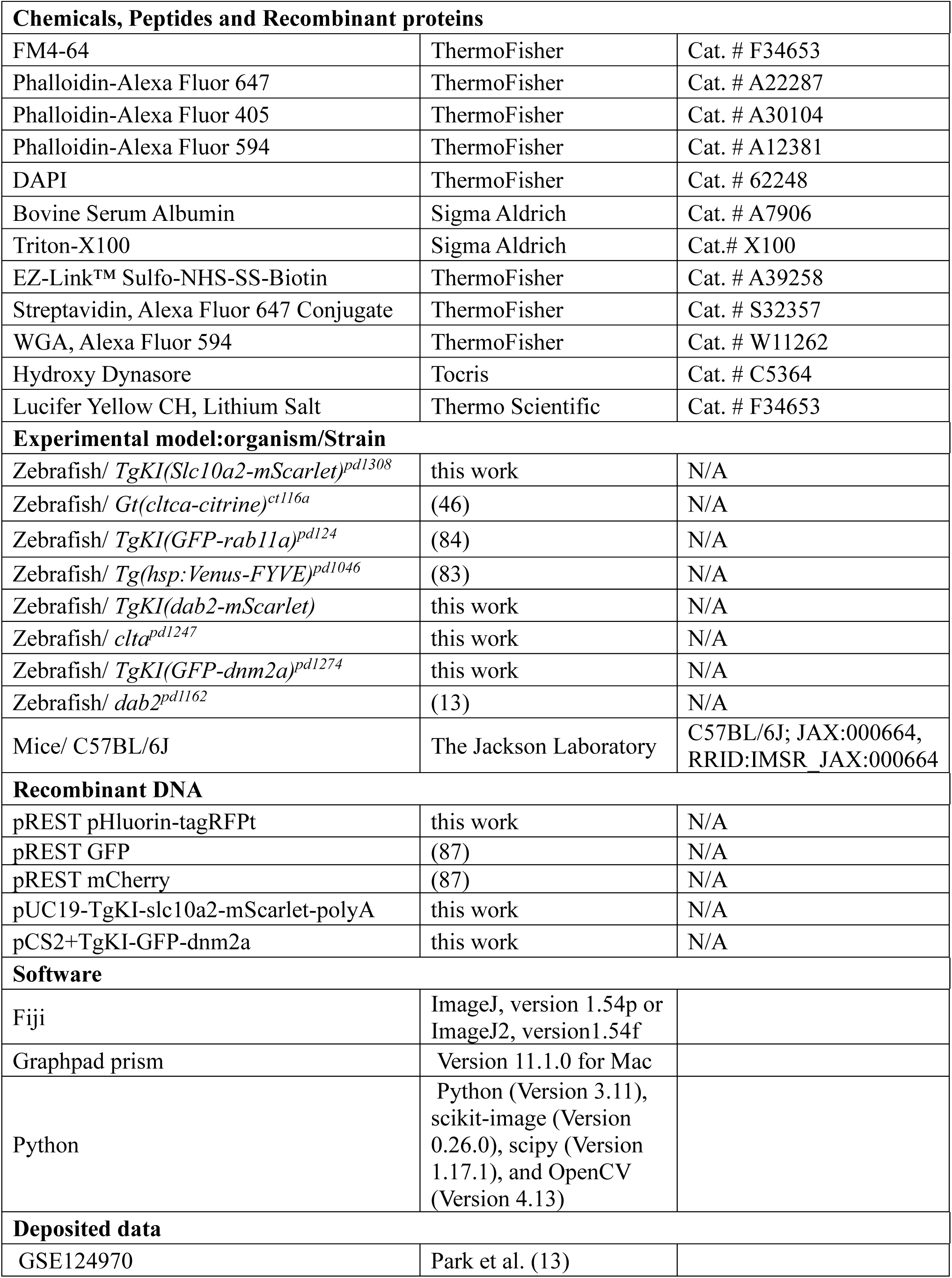

---

**Fig. S1. Related to Fig. 1.**
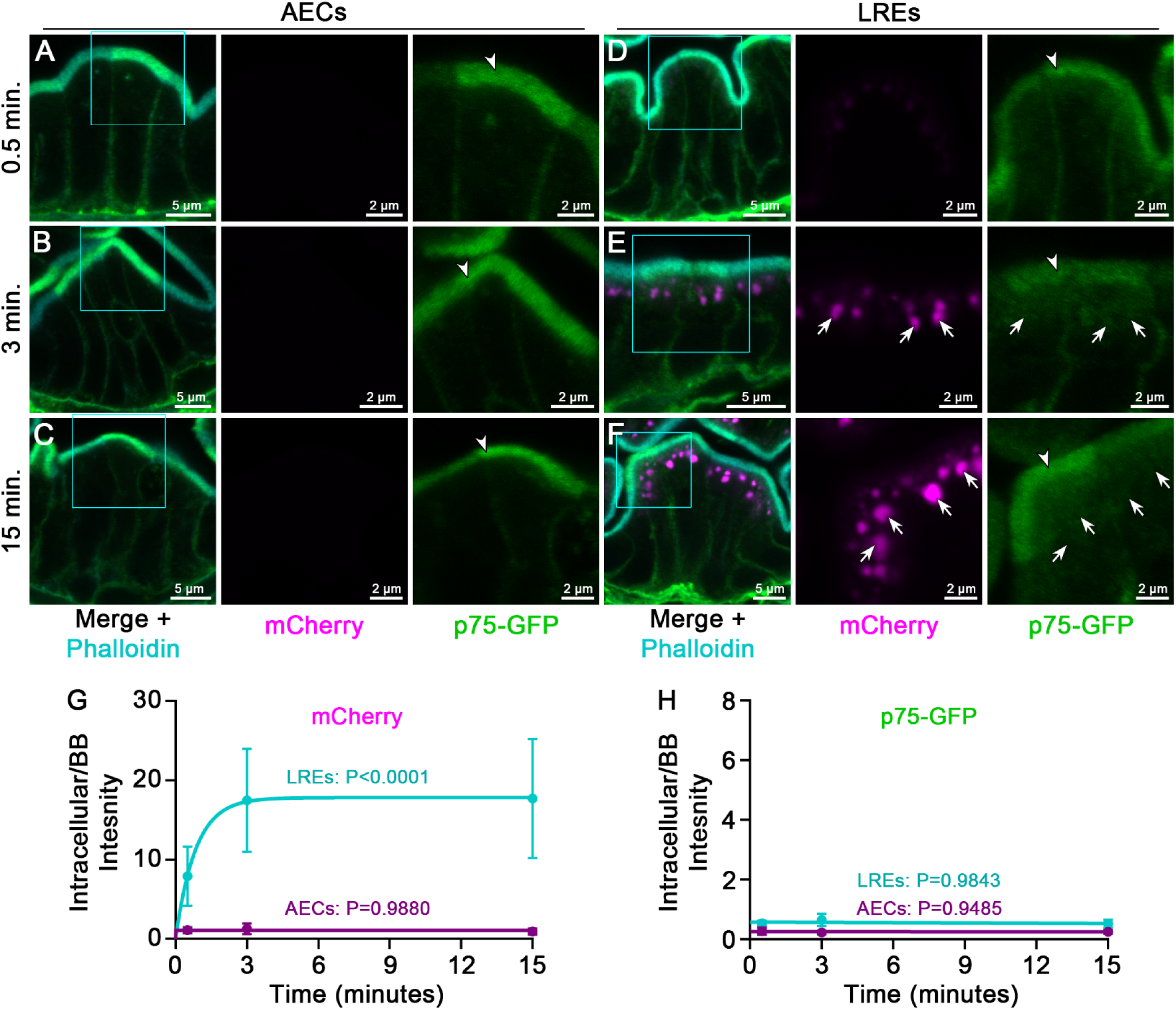
**(A-F)** Confocal images of AECs (A, B, C) and LREs (D, E, F) of 6 dpf *Tg(hsp70l:p75-GFP)^pd1180^* zebrafish larvae gavaged with mCherry (Magenta), fixed by 4% PFA gavage in 0.5 minutes (A, D), 3 minutes (B, E) and 15 minutes (C, F) following mCherry gavage and stained with Phalloidin-AF647 (F-actin, Cyan). Apical BB is marked by dashed lines using the Phalloidin-AF647 signal. White arrows show internalized mCherry (E, F) whereas white arrowheads show BB localization of p75-GFP (A-F). **(G-H)** Quantitation of Intracellular / BB intensity of mCherry (G) and p75-GFP (H) in LREs and AECs plotted against time (minutes). n ≥ 26 cells from ≥ 2 larvae. P values represent comparisons of t = 0.5 minutes versus t = 3 minutes and were calculated using Two-way ANOVA. Error bars are SD.

**Fig. S2. Related to Fig. 4.**
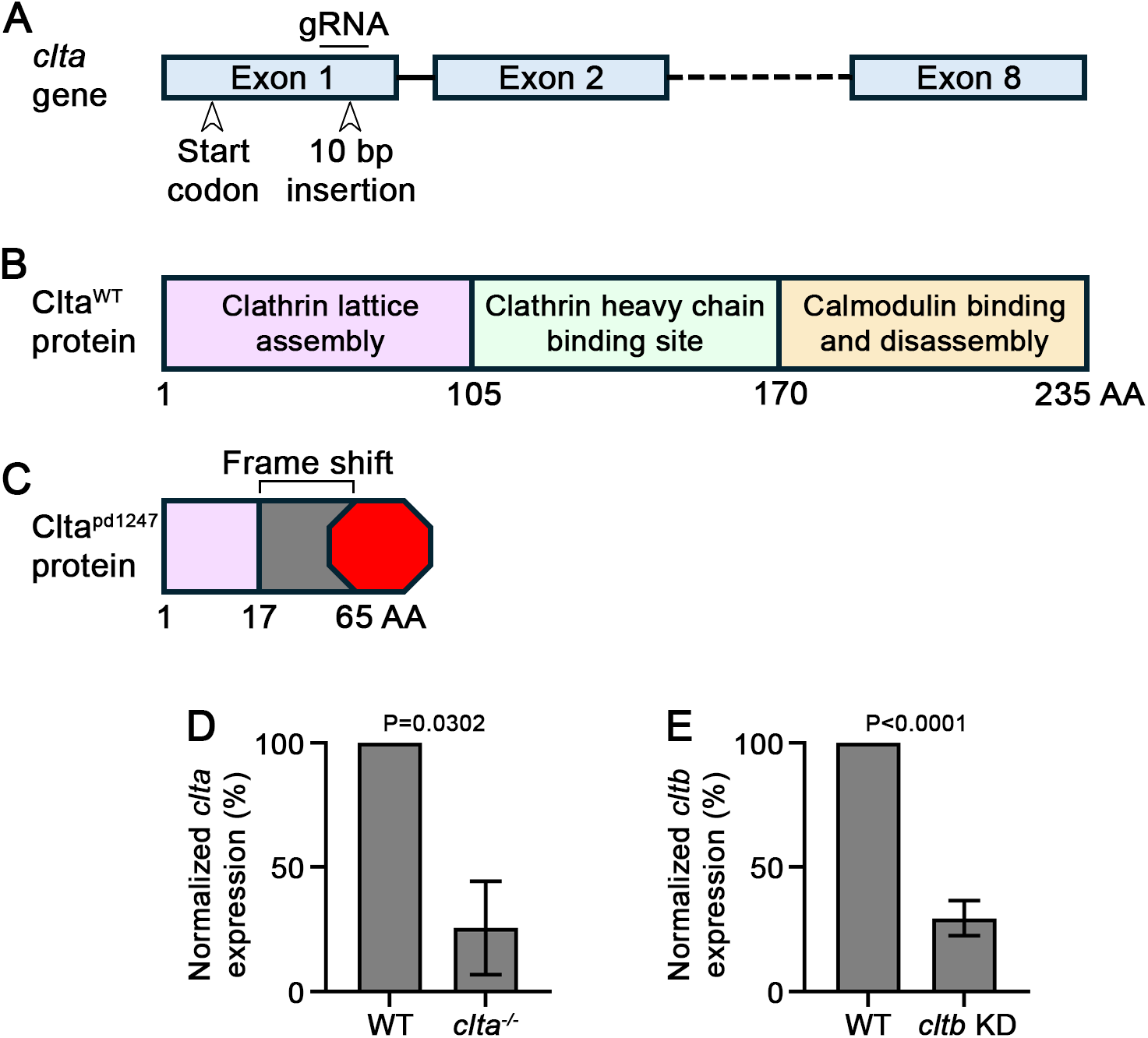
**(A-C)** Schematic diagram showing the zebrafish *clta* gene and gRNA target site for CRISPR mutagenesis (A), WT (B) and Clta^pd1247^ mutant protein (C). **(D)** Normalized *clta* mRNA expression in WT sibling and *clta^pd1247^* mutant zebrafish larvae by RT-PCR at 6 dpf. n = 2 biological replicates. **(E)** Normalized *cltb* mRNA expression in non-injected control and *cltb* CRISPR gRNA pool injected zebrafish larvae by RT-PCR at 6 dpf. n = 3 biological replicates. For panels D and E, error bars are SD and P values were calculated using an unpaired t test.

**Fig. S3. Related to Fig. 5.**
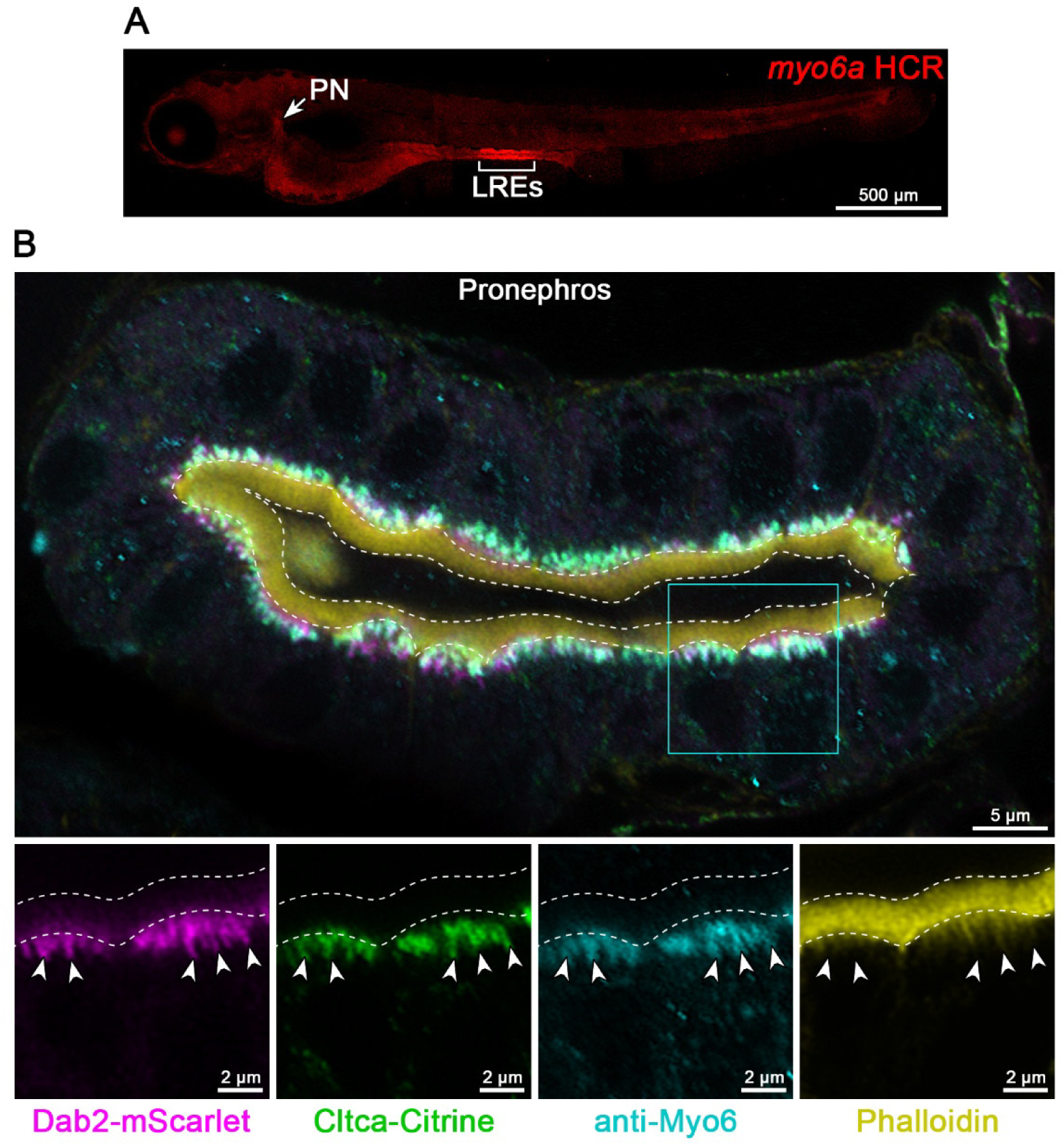
**(A)** Maximum intensity projections of a 6 dpf WT larva processed in situ hybridization chain reaction (HCR) detection of *myo6a* mRNA. Bracket indicates LREs, and arrow shows pronephros. **(B)** Confocal images of the pronephros region of 6 dpf *Gt(cltca-citrine)^ct116a^*; *TgKI(dab2-mScarlet)^pd1292^* larvae stained with anti-Myo6 antibody (Cyan) and Phalloidin-AF647 (yellow), showing co-localization of Myo6 (Cyan) and Dab2-mScarlet (Magenta) with Cltca-citrine-labeled ET (Green) by white arrowheads. Cyan box is the ROI for magnified insets.

**Fig. S4. Related to Fig. 6.**
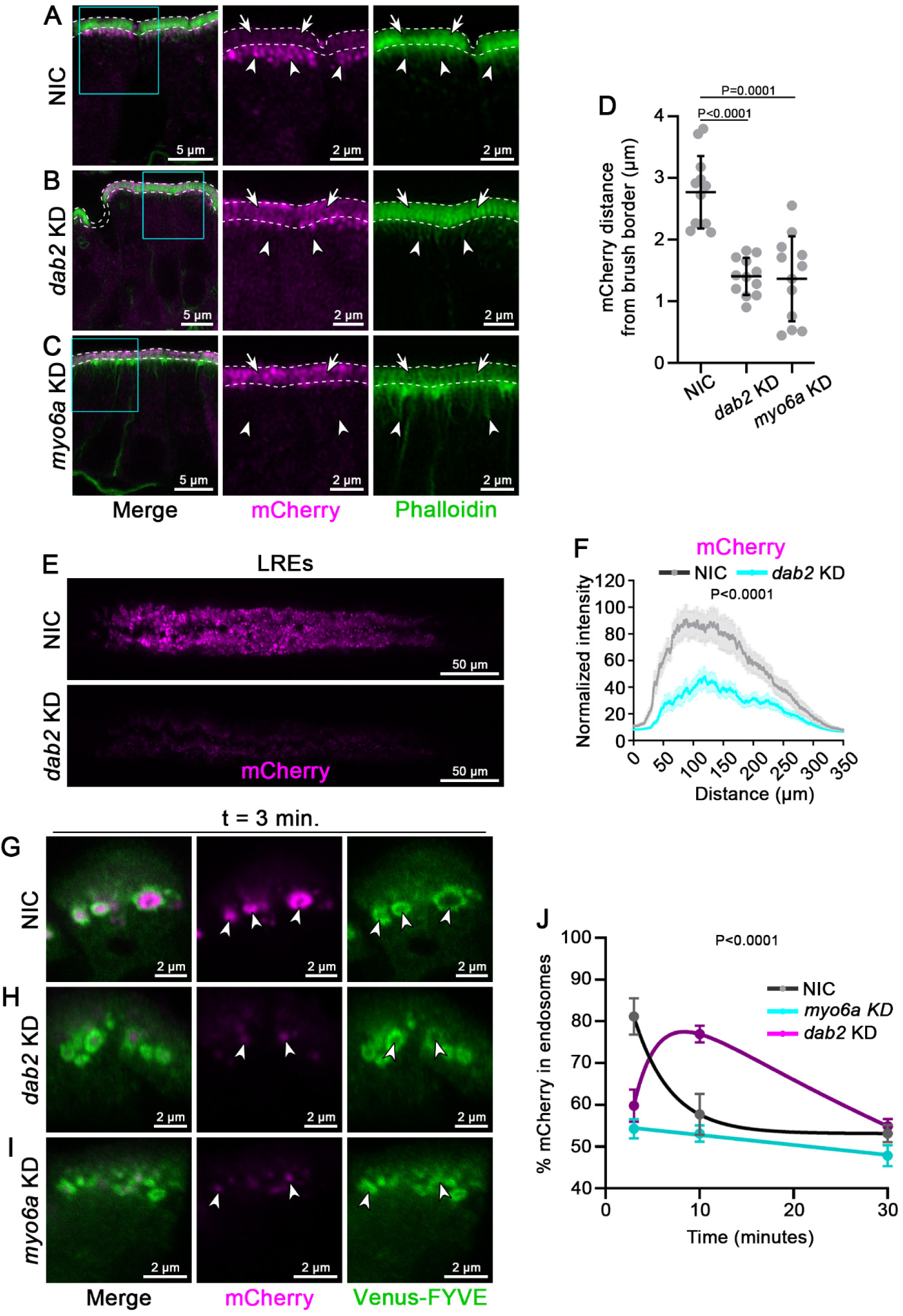
**(A-C)** Confocal images of 6 dpf NIC (A), *dab2* KD (B), and *myo6a* KD (C) of larvae gavaged with 4% PFA to block endocytosis followed by mCherry (Magenta) gavage, post-fixation, and staining with Phalloidin-AF647 (F-actin, Green). BB is marked by dashed lines. White arrows point to BB localization, and white arrowheads point to ETs (A) and corresponding sub-apical localization (B, C). Cyan boxes are ROIs for magnified insets. **(D)** Quantitation showing relative distribution of mCherry relative to the BB in NIC, *dab2* KD, and *myo6a* KD of 6 dpf larvae from panels A-C. Data points are distances between mean peak of mCherry intensity profile and BB tip within the ROI represented by red rectangle for n = 12 ROIs across n = 3 animals per group. Error bars are SD. P values were calculated using One-way ANOVA. **(E)** Maximum intensity projections of LRE regions showing mCherry uptake in 6 dpf NIC (top) and *dab2* KD larvae (bottom) 4 hours after gavage. **(F)** Quantification of total mCherry uptake for panel E. n ≥ 16 larvae per group. Error bars are SEM. P value was calculated using Two-way ANOVA. **(G-I)** Confocal images of NIC (G), *dab2* KD (H), and *myo6a* KD (I) LREs from 6 dpf *Tg(hsp:Venus-FYVE)^pd1046^* larvae gavaged with mCherry (Magenta) and fixed by 4% PFA gavage 3 minutes later, showing localization of luminal cargo with respect to Venus-FYVE positive early endosomes (Green) (white arrowheads). **(J)** Percentage of total internalized mCherry in endosomes marked by Venus-FYVE in LREs of 6 dpf *Tg(hsp:Venus-FYVE)^pd1046^*zebrafish larvae gavaged with mCherry (Magenta) and fixed by 4% PFA gavage at 3, 10, and 30 minutes after mCherry gavage. NIC: n = 8 cells from 6 larvae, t = 3 minutes; n = 9 cells from 4 larvae, t = 10 minutes; n = 11 cells from 5 larvae, t = 30 minutes. *dab2* KD: n = 7 cells from larvae, t = 3 minutes; n = 11 cells from 4 larvae, t = 10 minutes; n = 15 cells from 6 larvae, t = 30 minutes. *myo6a* KD: n = 10 cells from 4 larvae, t = 3 minutes; n = 8 cells from 3 larvae, t = 10 minutes; n = 4 cells from 2 larvae, t = 30 minutes. Error bars are SEM. P value was calculated using Two-way ANOVA and is a multiple comparisons test for NIC versus both *dab2* KD and *myo6* KD at t = 3 minutes.

## Notes

### Competing Interest Statement

The authors have declared no competing interest.

### Summary of Updates

We have removed the model in Fig. 6 and replaced this with a graphical abstract, added the word "endocytic" to the title, and slightly modified one sentence in the abstract. We also corrected minor errors in the figure legend for Fig. 3Z and Fig. 4T.

